# Identification of a stress-sensitive anorexigenic neurocircuit from medial prefrontal cortex to lateral hypothalamus

**DOI:** 10.1101/2021.09.07.459350

**Authors:** Rachel E Clarke, Katharina Voigt, Alex Reichenbach, Romana Stark, Urvi Bharania, Harry Dempsey, Sarah H Lockie, Mathieu Mequinion, Moyra Lemus, Bowen Wei, Felicia Reed, Sasha Rawlinson, Juan Nunez-Iglesias, Claire J. Foldi, Alexxai V. Kravitz, Antonio Verdejo-Garcia, Zane B. Andrews

**Affiliations:** Monash Biomedicine Discovery Institute and Department of Physiology, Monash University, Clayton, Victoria, 3800, Australia; Turner Institute for Brain and Mental Health, Monash University, Clayton, Victoria, 3800, Australia; Departments of Psychiatry, Anesthesiology, and Neuroscience, Washington University in St Louis, St Louis, MO, US; Monash Biomedicine Discovery Institute and Department of Anatomy and Developmental Biology, Monash University, Clayton 3800, Victoria, Australia

## Abstract

By modeling neural network dynamics related to homeostatic state and BMI, we identified a novel pathway projecting from the medial prefrontal cortex (mPFC) to the lateral hypothalamus (LH) in humans. We then assessed the physiological role and dissected the function of this mPFC-LH circuit in mice. *In vivo* recordings of population calcium activity revealed that this glutamatergic mPFC-LH pathway is activated in response to acute stressors and inhibited during food consumption, suggesting a role in stress-related control over food intake. Consistent with this role, inhibition of this circuit increased feeding and sucrose seeking during mild stressors, but not under non-stressful conditions. Finally, chemogenetic or optogenetic activation of the mPFC-LH pathway is sufficient to suppress food intake and sucrose-seeking in mice. These studies identify a glutamatergic mPFC-LH as a novel stress-sensitive anorexigenic neural pathway involved in the cortical control of food intake.

## Introduction

Appetite regulation is driven by genes implicated in the neuronal control of feeding (1, 2). Significant advances using animal models have revealed specific neural circuits controlling aspects of appetite, body weight control and energy balance, although these have largely centered on homeostatic systems governed by the hypothalamus and brain stem (3). A modern environment poses several challenges for biological systems to regulate food intake and body weight, such as the need to encode and balance physiological interoceptive signals in combination with motivational and affective inputs, including reward and stress. In line with this, attention has now shifted to the higher-order cognitive and emotional processes that influence feeding behavior, encoded by corticolimbic neural structures (4, 5).

The medial prefrontal cortex (mPFC) is consistently identified as a key brain region implicated in the valuation of food (6, 7). mPFC activity is associated with subjective health and taste ratings of food in self-reported dieters, and this activity varies based on whether an individual exerts dietary self-control or not (7). In addition, mPFC activity is greater when individuals are asked to choose between healthy and unhealthy food options in a fasted state relative to a sated state (6), suggesting the mPFC is sensitive to homeostatic information. People with obesity consistently display stronger activation of the mPFC in response to images of food than healthy weight controls (8–10). However, it is not possible to conclude that greater activity of the mPFC drives overconsumption of palatable food. For example, studies in patients with anorexia nervosa also show similar elevated mPFC activity in response to images of foods compared to healthy controls (11). Thus, it is difficult to accurately assign a function based on brain imaging of a single region, reinforcing the notion that diverse connectivity patterns to downstream structures must be examined (12).

Human imaging studies demonstrate functional connectivity between the hypothalamus and a broad range of cortical and subcortical structures, including those involved in food valuation (13, 14). More specifically our recent studies demonstrated that the mPFC is functionally connected with the lateral hypothalamus (6), a brain region well known to regulate food intake, motivation and reward (15). However, functional connectivity assessed through fMRI does not directly represent anatomical connectivity or functionality, as several regions that are not connected still exhibit functional correlations (16). To gain greater insight into the relationship between the mPFC and hypothalamus, we applied a novel neural network modelling approach to previously collected data across a wide BMI range under both hunger and sated conditions (6). Results from this analysis predicted the presence of a specific neural circuit from the mPFC to the lateral hypothalamus (LH), which was differentially affected by hunger. We then sought to test the function of this circuit using mouse models that allow for specific manipulation of the mPFC-LH circuit *in vivo*.

Although anatomical mPFC projections to the LH have been known for decades (17, 18), the function of this circuit has received very little attention in animal research. Studies suggests a role for the mPFC-LH circuit in aggressive behavior in response to changing social threats (19); impulsive responding in a go/no task (20); and promotion of social dominance (21). Whether or not the mPFC-LH affects food intake and appetitive behavior has not yet been thoroughly addressed. Therefore, we examined the functional role of this mPFC-LH circuit in feeding and motivated behavior in mice using fiber photometry, chemogenetics and optogenetics. Our results show that the mPFC-LH pathway is activated by acutely stressful stimuli and inhibited by food consumption, independent from food palatability or metabolic state. As predicted from human neural network modelling, we show that activation of the mPFC-LH pathway reduces feeding and food motivation, whereas acute inhibition promotes food intake and motivation in stressful environments. Thus, we have identified a novel stress-sensitive anorexigenic pathway from the mPFC to the LH.

## Methods and Materials

### Spectral Dynamic Causal Modeling of network function

Participants completed two resting-state fMRI scans, one after an overnight fast (hunger condition) and one after a standardized breakfast (satiety condition), as previously described and reported (6). Briefly, participants were instructed to have a typical meal (700-1000kj) between 7.30pm and 8.30pm on the night prior to their scan and refrain from eating or drinking (except for water) until their morning scan. For the satiety condition, participants received a standardized breakfast (293 kcal) one hour prior to their scan. Data collected from functionally connected brain regions during resting state fMRI scans were then used to model network function through spectral dynamic causal modeling. To address our main hypotheses, we focused on spectral dynamic causal modeling analyses that assessed changes in effective connectivity of fasting versus satiety condition independent of BMI (main effect of fasting), changes in effective connectivity modulated by BMI (main effect of BMI), and changes in fasting-related effective connectivity modulated by BMI (BMI-by-fasting interaction). Importantly, spectral dynamic causal modelling predicts effective connectivity, which is the influence that one brain node exerts over another through a network model of causal dynamics. This approach requires input data from functionally connected brain regions identified in fMRI studies and thus, the output data represent a prediction about the direction and valence of an identified pathway using the defined experimental parameters. For further details see Supplemental Methods.

### Animal Studies

All experiments were conducted in accordance with the Monash Animal Ethics Committee guidelines. C57BL/6 male mice were obtained from the Monash Animal Services facility. Vglut1-ires-cre male mice on a C57BL/6 background were obtained from the Jackson Laboratory (B6.Slc17a7-IRES2-Cre-D; stock number 023527) and bred in the Monash Animal Services Facility. All mice were aged between 8-10 weeks at beginning of experiments. Mice were maintained on a 12-hour light-dark cycle with *ad libitum* access to standard chow (Specialty Feeds, Western Australia) and water under standard laboratory conditions (21°C).

### Viruses and Surgical procedures

Surgical procedures and virus details can be found in supplemental methods and materials.

### Behavioral Testing

For detailed description of behavior experiments and operant conditioning experiments using the feeding experimental devices 3 (FED3; Open Ephys, Portugal) (22) see supplemental methods and materials.

### Fiber photometry recording and analysis

A detailed description of the fiber photometry setup, behavioral experiments and analysis can be found in the supplemental methods and materials. Data analysis for each experiment was performed using a custom python code (https://github.com/Andrews-Lab/Clarke-et-al-mPFC-LH-manuscript).

### Statistical Analysis

Data are expressed as mean +/- standard error of the mean. Comparisons were tested using two-tailed unpaired t-tests or paired t-tests. A one-way ANOVA with Tukey’s multiple comparison test was applied when multiple groups were compared at the same time point. A repeated measures one-way ANOVA was applied when multiple time points were compared to baseline values. A two-way ANOVA with Sidak’s multiple comparison test was applied when both fasting and fed conditions were present, or when a variable was measured over time. All statistical tests used are described in figure legends with details provided in supplementary table 2.

## Results

### Identification and functional dissection of the mPFC-LH pathway

We used the time-series from the cortical and hypothalamic regions of interest from previously published fMRI data to define and estimate a fully connected dynamic causal model for each participant and condition using Variational Laplace (23) (Fig S1). Analysis of these models revealed that under conditions of hunger, there is reduced inhibitory influence from the ventral medial prefrontal cortex (vmPFC) bilaterally to the LH (S1), suggesting that this pathway may normally restrain feeding or represent a coping mechanism to hunger. This assessment of the effective connectivity, however, cannot mechanistically test the biological function of a predicted pathway from the mPFC-LH. Here, we have used preclinical models, coupled with neural circuit recordings and manipulations to ascertain the function of mPFC-LH pathway. In this manner, we have leveraged human neural network modeling to inform and guide our preclinical studies.

To begin the preclinical mouse studies, we first confirmed the presence of monosynaptic mPFC to LH projections in mice. We injected a retrogradely transported viral vector expressing GFP (AAVrg-CAG-GFP), bilaterally into the LH (Fig 1A, B) and identified cell bodies in the mouse prelimbic, infralimbic and rostral cingulate gyrus 1 (Cg1) regions (Fig 1C), which we refer to as mPFC, as described previously (24, 25). For continuity with our human data described above, we refer to projections from these regions to the LH as mPFC–LH. We also observed cells in the insular cortex (Fig 1C), consistent with a known LH input from the insular cortex (26).

**Figure 1.**
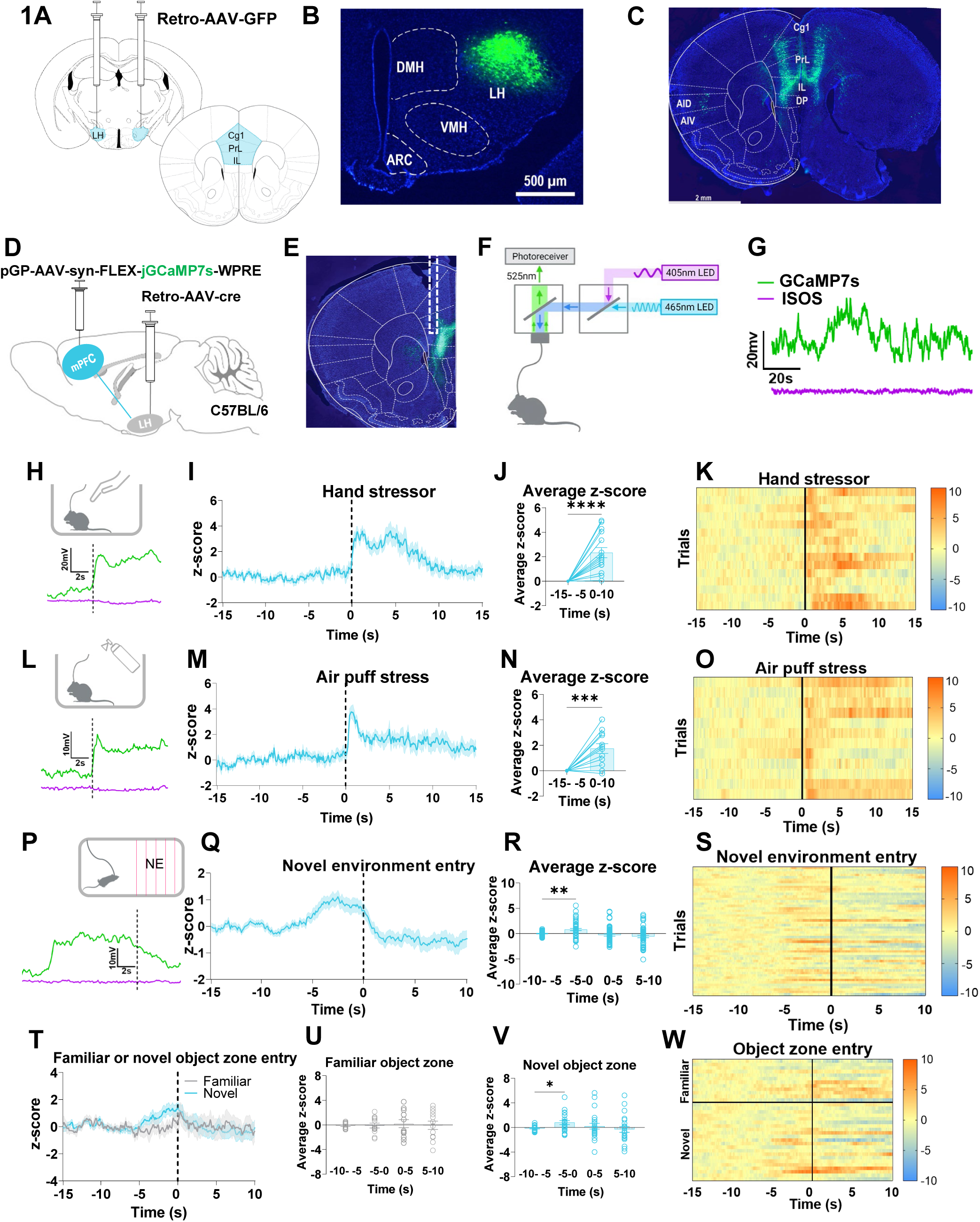
mPFC-LH projecting neurons respond to acute stressors and novelty. **A** Schematic illustrating lateral hypothalamus (LH) target sites for retrograde tracing (left) and the regions of the mouse medial pre-frontal cortex (mPFC) where projecting neurons are observed (right): Cg1, Cingulate gyrus 1; PrL, Prelimbic cortex; and IL, Infralimbic cortex. **B** Representative coronal section of LH following injection of AAVrg-CAG-GFP (green) with DAPI staining (blue). DMH, Dorsomedial hypothalamus; VMH, Ventral medial hypothalamus; ARC, Arcuate nucleus of the hypothalamus. **C** Representative coronal section showing expression of GFP (green), with DAPI staining (blue), throughout the Cg1, PrL and IL regions of the mPFC following bilateral injection of AAVrg-CAG-GFP into the LH. AID, Agranular insular cortex dorsal part; AIV, Agranular insular cortex ventral part; DP, Dorsal peduncular cortex, Cg1, Cingulate gyrus 1; DP, Dorsal peduncular cortex; fmi, forceps minor of the corpus callosum, IL, Infralimbic cortex; and PrL, Prelimbic cortex. **D** Schematic illustrating the viral approach used to record from mPFC-LH projecting neurons. **E** Representative image showing GcaMP7s expression in mPFC-LH projecting neurons and fiber optic cannula placement. **F** Fiber photometry approach used to monitor neural activity in mPFC-LH projecting neurons with 405 nm light used as an isosbestic control for motion artefact, created with BioRender.com. **G** Short trace demonstrating calcium-dependent fluorescence associated with GcaMP7s (465 nm) and calcium-independent fluorescence associated with the isosbestic control (405 nm). **H** Hand in cage experiments with example trace (n=5 mice, n=3 trials per animal, 15 trials in total). **I** mPFC-LH projecting neurons are acutely activated by a hand in cage stressor (Video 1). **J** Average z-score was significantly greater 0-10 seconds after hand entered cage compared to -15- -5 seconds (baseline period) prior to hand in cage (two-tailed paired t-test, p<0.0001). **K** Heat maps showing individual trial responses to a hand in cage test in order of trial. **L** Air puff stressor with example trace (n=4 mice, n=3 trials per animal, 12 trials in total). **M** mPFC-LH projecting neurons are acutely activated by an air puff stressor (Video 2). **N** Average z-score was significantly greater 0-10 seconds after hand entered cage compared to -15- -5 seconds (baseline period) prior to hand in cage (two-tailed paired t-test, p=0.0009). **O** Heat maps showing individual trial responses to air puff in order of trial. **P** Novel environment stressor with example trace (n=6 mice, n=9 entries per animal, 54 trials in total). **Q** mPFC-LH projecting neuronal activity increases prior to voluntary entry to a novel environment. **R** Average z-score is significantly greater -5-0s before entering the novel arena compared to - 10- -5s (baseline period) before entry (repeated measures one-way ANOVA with dunnet’s multiple comparison test, -5 to 0s>-10- -5s, p=0.0015). **S** Heat maps showing individual trials of novel environment entry in order of trial. **T** mPFC-LH activity during entry into a familiar object zone (grey) or a novel object zone (blue) (n=4 mice, 2-6 zone entries). **U** Average z-score following entry into the familiar object zone did not differ significantly from baseline at - 5-0s before entry or 0-5s and 5-10s following entry. **V** Average z-score was significantly higher -5-0s prior to entry into the novel object zone than baseline (one-way ANOVA with Dunnet’s multiple comparison test, -5-0>-10 -5s, p=0.0171). **W** Heatmap of individual trials into the familiar and novel object zones in order of trial with example traces from a familiar entry (top left) and novel entry (bottom left). All data are mean +/- SEM. Circles represent individual trials. Dashed lines at 0 (**I**, **M**, **Q**, **T**) indicate time at which stimulus or event occurred. *p<0.05, * p<0.01, ***p<0.001, ****p<0.0001.

### Acute stressors modulate the mPFC-LH circuit

To identify how the mPFC-LH pathway is physiologically modulated, we used *in vivo* fiber photometry coupled with our dual viral approach (Fig 1D-G). The potential anorexigenic nature of this circuit suggested by human network modeling, indicated it may be activated by factors that attenuate feeding such as acute stressors. To test this, mice were exposed to hand in cage stress and air puff stress (Supplementary videos 1&2). Both hand in cage and air puff stress increased mPFC-LH neural activity, as measured by GCaMP7s fluorescence, resulting in an approximately 2-fold increase in the first 10 seconds after stress exposure relative to the baseline period (−15 to −5 seconds prior to stress exposure) (Fig 1H-O; Videos 1 and 2). The increased activity occurs with the appearance of the hand and peaks at contact with the hand (Fig S2D-E), similar in profile to stress-sensitive corticotropin-releasing hormone neurons (27). Novel environments are known to produce acute anxiety-like behavior (28) and we examined the impact of a novel environment on mPFC-LH activity. Although mPFC-LH activity increased immediately prior to entry into the novel environment, it did not vary significantly from baseline following entry into the novel environment (Fig 1P-S). Moreover, we noticed a significantly higher peak z-score when comparing the 2^nd^ entry to the last entry, suggesting that neural responses were reduced as animals became more familiar with the environment (Fig S2F-G). Video analysis of photometry sessions revealed that mice made significantly more entries into the novel environment, and the velocity within the novel environment was significantly greater compared to the familiar environment (Fig S2H-K). Similarly, we observed a greater increase in mPFC-LH activity when mice approached a novel object versus a familiar object (Fig 1T-W). Collectively, these results suggest that mPFC-LH activity is responsive to acute stressors and is transiently elevated prior to interaction with novel environments and stimuli.

### Food consumption inhibits the mPFC-LH circuit

We then exposed mice to chow, familiar palatable or novel palatable foods and examined the effects on mPFC-LH activity. When mPFC-LH activity was aligned to food consumption, we observed a significant reduction in response to chow, familiar palatable and novel palatable foods but not in response to chewing a wooden dowel (Fig 2A-C). Moreover, the change in activity occurred upon consumption and not in response to presentation of the food or wooden dowel or upon approach (Fig S3A-B). This change was not proportional to calories consumed, palatability of food (Fig 2D-E), or metabolic state (Fig S3A-G), suggesting this circuit is not part of a broader neural network responding to calorie content or metabolic need.

**Figure 2.**
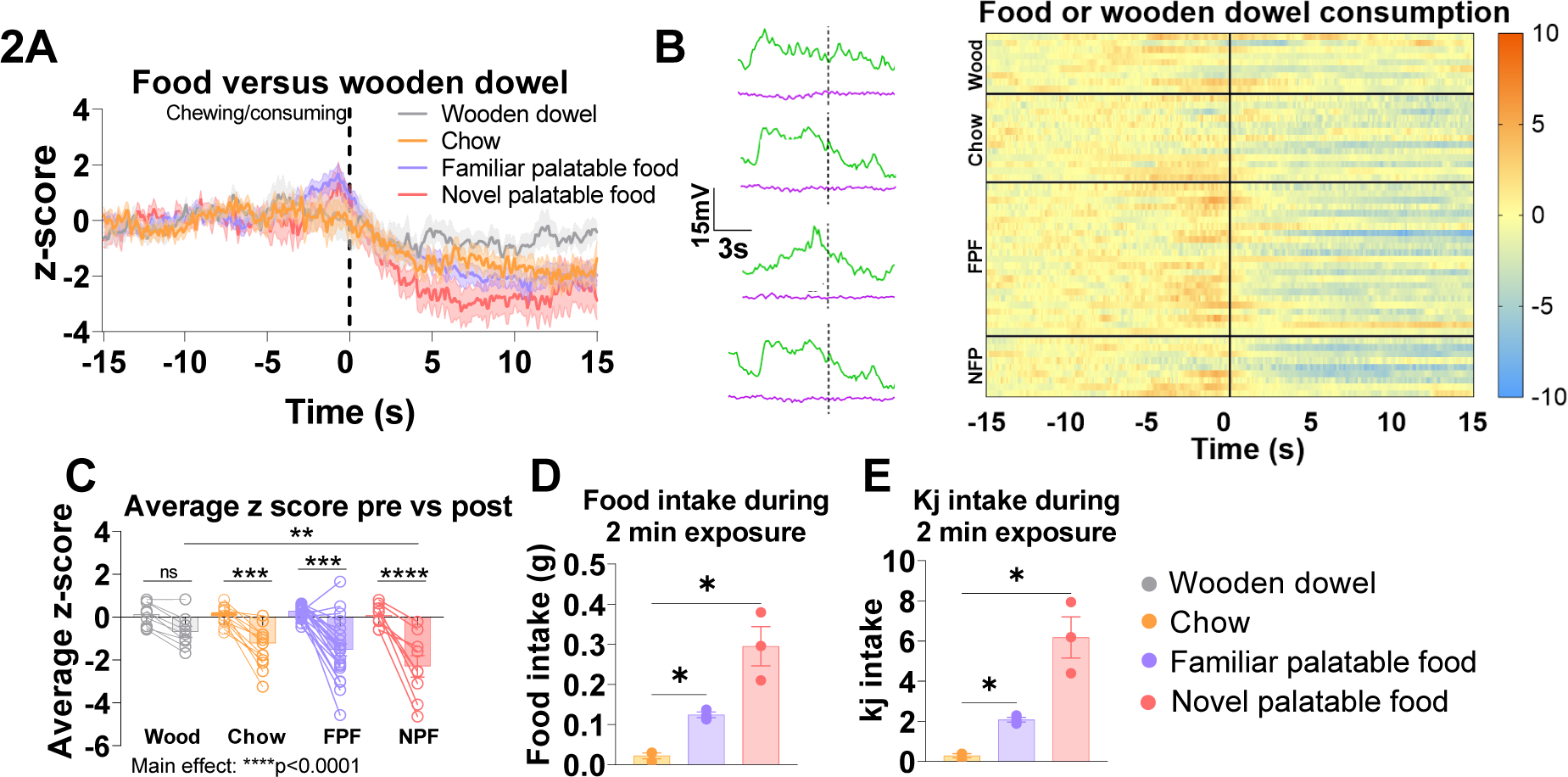
mPFC-LH circuit is suppressed during food consumption but not chewing of a non-food object. **A** mPFC-LH projecting neuron activity during chewing and consumption of wooden dowel (n=3 mice, n=1-4 trials per animal, 9 trials in total, grey); chow (n=3 mice, n=2-8 trials per animal, 13 trials in total, orange); familiar palatable food (FPF) (n=3 mice, n=5-10 trials per animal, 23 trials in total, purple); and novel palatable food (NPF) (n=3 mice, n=2-4 trials per animal, 9 trials in total, pink) **B** Heatmap showing individual trials for chewing and consumption of wooden dowel, chow, familiar palatable food and novel palatable food in order of trial with example traces for each experiment (left). **C** Average z-score 0-15s post chewing wooden dowel did not differ from -15-0s pre chewing. Average z-score 0-15s post chewing and consumption of chow, familiar palatable food and novel palatable food was significantly lower than -15-0s pre consumption (ordinary one-way ANOVA with planned comparisons and Sidak’s multiple comparison test: chow -15-0s>0-15s p=0.0007; familiar palatable food -15-0s>0-15s p<0.0001; novel palatable food -15-0s>0-15s p<0.0001). Average z-score 0-15s post consumption of a novel palatable food is significantly lower than average z-score 0-15s post chewing and consumption of wooden dowel (wooden dowel 0-15s > novel palatable food 0-15s, p=0.003). **D** Food intake during the 2-minute exposure to chow (orange), familiar palatable food (purple), and novel palatable food (pink) (n=3 mice, average over 2-10 trials). Mice ate significantly more familiar palatable food and significantly more novel palatable food compared to chow (repeated measures one-way ANOVA with Tukey’s multiple comparison test; familiar novel food>chow, p=0.0339; novel palatable food>chow, p=0.0423). **E** Mice consumed significantly more kilojoules (Kj) from the familiar palatable food and novel palatable food compared to chow (repeated measures one-way ANOVA with Tukey’s multiple comparison test; familiar novel food>chow, p=0.0244; novel palatable food>chow, p=0.0443). All data are mean +/- SEM. Circles represent individual trials. Dashed lines at 0 (**A**) and solid lines at 0 (**B**) indicate time at which chewing or consumption occurred. *p<0.05, *p<0.01, ***p<0.001, ****p<0.0001. LH, lateral hypothalamus; mPFC, medial pre-frontal cortex; ns, not significant

### Inhibition of the mPFC-LH pathway increases food intake

To test if inhibition of the mPFC-LH pathway could increase food consumption, we expressed the inhibitory DREADD receptor hM4Di in mPFC-LH projecting neurons (Figure 3A). Cre-dependent mCherry and hM4Di expression was observed in the mPFC, and fos activity was significantly reduced in hM4Di-expressing animals compared to mCherry controls following CNO (Fig 3B-C). Inhibition of the mPFC-LH pathway did not affect food intake in a non-stressed home cage environment in the light phase (Fig 3D, G), after an acute 3 hour fast at the beginning of the dark phase (Fig 3E, H) or after a 14-hour fast (Fig 3F, I). Similarly, motivated home cage food-seeking as assessed with a progressive ratio with FED3 showed no differences in breakpoint reached during either fed or fasted conditions (Fig 3J-K). Since our photometry data suggested the mPFC-LH pathway is activated by acute stress and novelty (Fig 1H-W), we reasoned that the lack of effect on home-cage feeding was due to already low mPFC-LH in a familiar environment.

**Figure 3.**
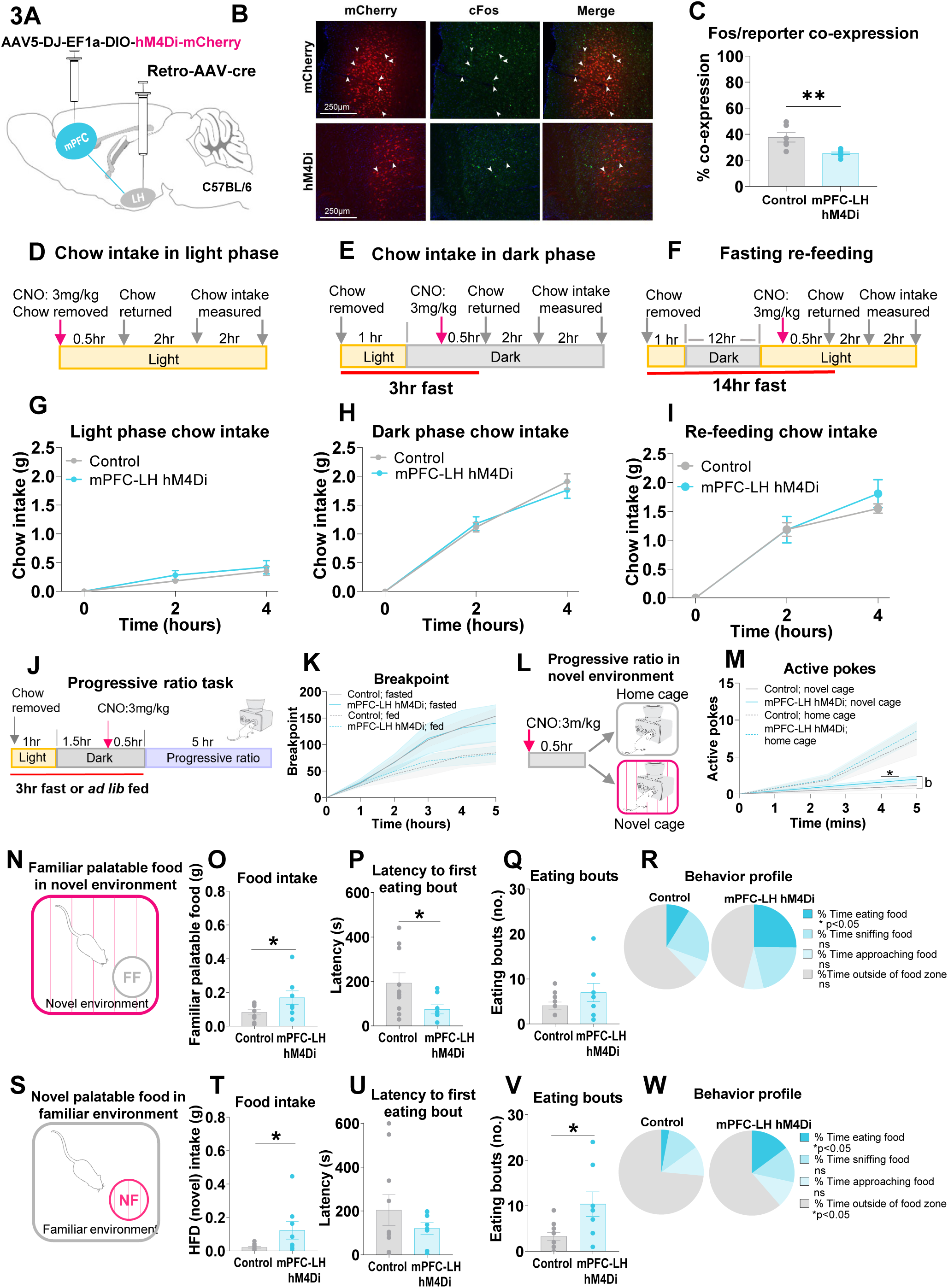
Chemogenetic inhibition of the mPFC-LH circuit promotes feeding and reward seeking in mildly stressful conditions. **A** Schematic illustrating the viral injection strategy for chemogenetic inhibition of mPFC-LH projecting neurons using hM4Di-mCherry DREADDs. Control mice received an injection of a cre-dependent virus encoding for the mCherry reporter protein only. **B** Representative coronal sections of mPFC collected 90 minutes after intraperitoneal injection (i.p.) of CNO (3mg/kg). Immunofluorescent staining shows mCherry (red) and c-Fos (green), a marker of neuronal activity with co-expression is indicated by white arrows **C** Percentage co-expression of reporter protein and Fos (n=6 for control, n=7 for mPFC-LH hM4Di, two-tailed unpaired t-test, p=0.0049). **D-F** Schematics illustrating the experimental timelines for food intake experiments performed under varying conditions and phases of the light-dark cycle. CNO (3mg/kg) was injected 30 mins prior to the beginning of each experiment (red arrow). **G-I** Inhibition of mPFC-LH projecting neurons had no effect on feeding in the home cage at any stage of the light-phase or following an overnight fast (n=7-11 for control, n=5-7 for mPFC-LH hM4Di, two-way ANOVA). **J** Schematic illustrating the experimental timeline for the progressive ratio task using the FED3. All mice were tested in an *ad lib* fed state and following a short fast in sessions separated by one week. Mice were counterbalanced to the order of treatment. See methods for details. **K** Inhibition of mPFC-LH projecting neurons in the home cage had no effect on breakpoint ratio in the progressive ratio task (n=6-9 for control, n=7 for mPFC-LH hM4Di, two-way ANOVA). **L** Schematic illustrating the experimental timeline for the progressive ratio in the novel environment and home cage, performed after the first progressive ratio task following a washout period. Mice were tested in the dark phase in both home cage and novel cage and were counterbalanced to the order of treatment. See methods for further details. **M** Inhibition of mPFC-LH projecting neurons had no effect on active pokes reached after a short progressive ratio task in the home cage (dashed lines, n=7 for control, n=6 for mPFC-LH hM4Di, two-way ANOVA), but did significantly increase the active pokes reached in the novel cage from 4 to 4.5 minutes (solid lines, n=7 for control, n=6 for mPFC-LH hM4Di, two-way ANOVA, p=0.0132) and resulted in a significant group-by-time interaction (p=0.0213). **N** Mice were placed in a novel environment with access to a familiar palatable food 30 minutes following CNO injection (n=10 for control, n=8 for mPFC-LH hM4Di, two-tailed unpaired t-test). **O** Inhibition of mPFC-LH projecting neurons significantly increased intake of the familiar palatable food compared to control (p=0.0457). **P** Latency to approach the familiar palatable was significantly reduced in the mPFC-LH hM4Di group (p=0.0439). **Q** Number of eating bouts were not significantly different between mPFC-LH hM4Di and control groups. **R** mPFC-LH hM4Di mice spent significantly more time eating than controls (p=0.0379), however time sniffing, approaching or time outside of the food zone did not vary between groups. **S** Mice were placed in a familiar environment with access to a novel palatable food 30 minutes following CNO injection (n=10 for control, n=8 for mPFC-LH hM4Di, two-tailed unpaired t-test). **T** Inhibition of mPFC-LH projecting neurons significantly increased intake of the novel palatable food compared to control (p=0.0477). **U** Latency to approach the novel palatable food was not significantly different between mPFC-LH hM4Di and control groups **V** Number of eating bouts was significantly increased with inhibition of mPFC-LH projecting neurons (p=0.0142). **W** mPFC-LH hM4Di mice spent significantly more time eating than controls (p=0.0197) and significantly less time outside of the food zone (p=0.0173), however time sniffing or approaching food zone did not vary between groups. All data are mean +/- SEM. For statistical analysis, a indicates a significant main effect of group, b indicates a significant interaction, *p<0.05 and **p<0.01 for between group comparisons. CNO, clozapine-N-Oxide; FED3, feeding experimentation device version 3; LH, lateral hypothalamus; mPFC, medial pre-frontal cortex; ns, not significant.

To examine the role of acute stress on motivated reward seeking, we conducted a progressive ratio experiment in a novel environment, a validated model of acute stress that affects feeding (29).Following CNO injections, we observed that active nose pokes were suppressed during the first 5 minutes compared to the home cage (Fig 3M), confirming that acute stress transiently affects motivated reward seeking. Intriguingly, inhibition of the mPFC-LH pathway with hM4Di attenuated this effect (Fig 3M), although significant differences were only seen at acute time points, indicating that the effect of novel environment stress wears off with time. Moreover, mPFC-LH inhibition promoted the consumption of familiar palatable food in a novel environment. (Fig 3N-R). We observed similar effects when novel palatable food was given in a familiar environment (Fig 3S-W). Although mPFC-LH inhibition produced no differences in anxiety-like behavior in the elevated zero maze, light-dark box, or open field (Fig S4), analysis of behaviors expressed in an open field revealed that mPFC-LH inhibition reduced time spent rearing (Fig S4M). Interestingly, rearing behavior is typically expressed within a novel environment to survey for potential danger or threat (30). Collectively, these results suggest that mPFC-LH inhibition alleviates the stress-induced decrease in motivation for sucrose pellets and the avoidance of novel food or food presented in a novel environment.

### Activation of the mPFC-LH pathway reduces food motivation

We then expressed the DREADD receptor hM3Dq to activate mPFC-LH projecting neurons (Fig 4A) and test the effect on food intake and motivation for sucrose pellets. This resulted in substantial expression of hM3Dq and mCherry in the mPFC with significantly greater Fos/mCherry co-expression after CNO (Fig 4B, C). First, activation of the mPFC-LH circuit during the early light phase, the period when mice are typically inactive (Fig 4D), significantly suppressed food intake after 4 hours (Fig 4G). We also examined food intake following a short-term 3hr fast to normalize feeding status during the dark phase, the period when mice consume most of their energy intake (Fig 4E), and again activation of the mPFC-LH circuit significantly suppressed food intake after 4 hours (Fig 4H). Finally, we investigated whether mPFC-LH circuit activation could suppress feeding when mice are challenged with a longer term overnight fast (Fig 4F) and found that circuit activation suppressed food intake at 4 hours (Fig 4I) from food being available. Together, these data show that activation of the mPFC-LH pathway is sufficient to suppress food intake, regardless of time of day or metabolic status.

**Figure 4.**
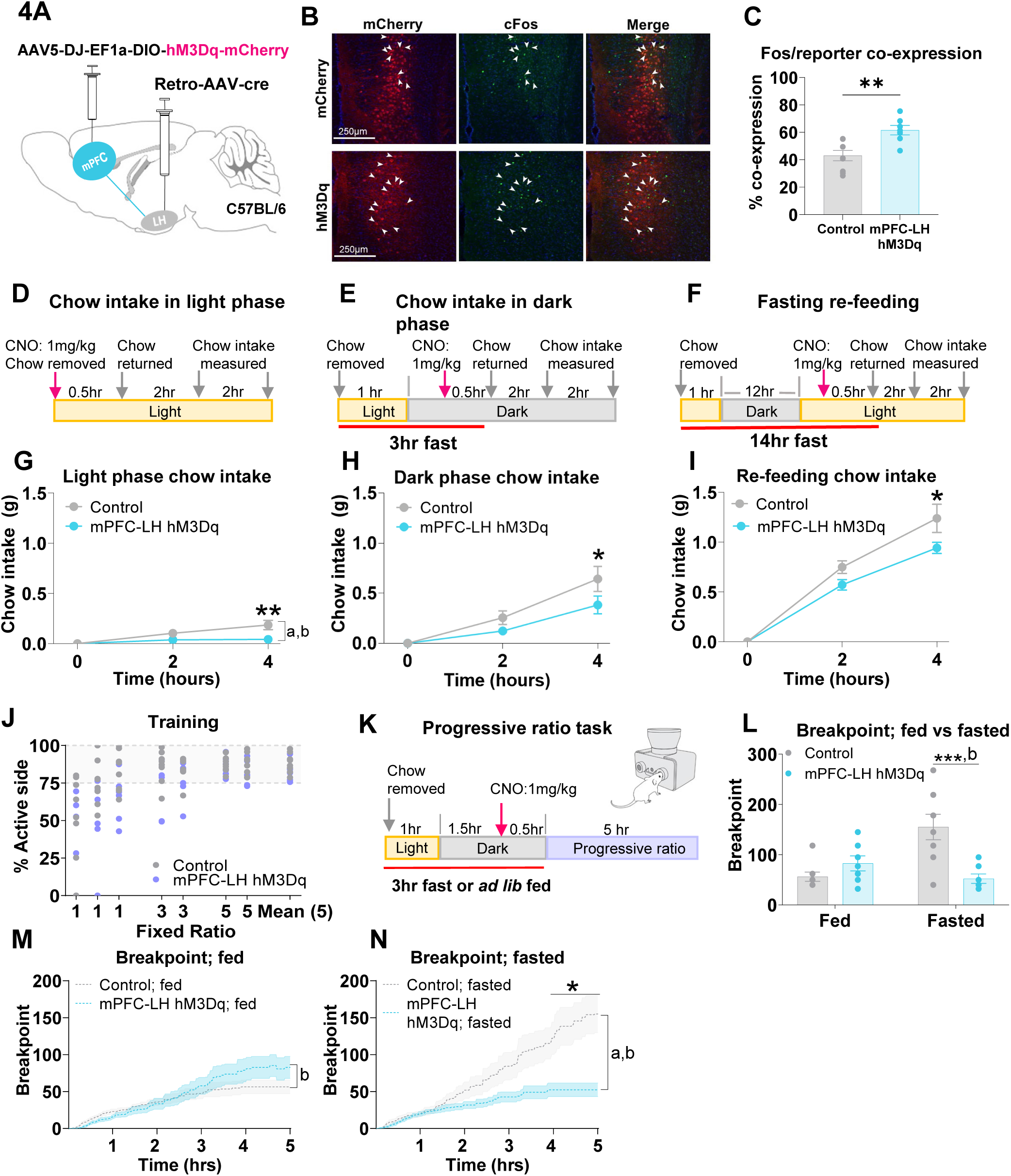
Chemogenetic activation of the mPFC-LH circuit in mice suppresses food intake and reward seeking. **A** Schematic illustrating the viral injection strategy for chemogenetic activation of mPFC-LH projecting neurons using hM3Dq-mCherry DREADDs. Control mice received an injection of a cre-dependent virus encoding for the mCherry reporter protein only. **B** Representative coronal sections of mPFC collected 90 minutes after intraperitoneal injection (i.p.) of CNO (1mg/kg). Immunofluorescent staining shows m-Cherry (red) and c-Fos (green), a marker of neuronal activity. Co-expression is indicated by white arrows. **C** Percentage co-expression of reporter protein and Fos (n=9 for control, n=7 for mPFC-LH hM3Dq, two-tailed unpaired t-test, p=0.0033). **D-F** Schematics illustrating the experimental timelines for food intake experiments performed under varying conditions and phases of the light-dark cycle. CNO (1mg/kg) was injected 30 mins prior to the beginning of each experiment (red arrow). **G-I** Activation of mPFC-LH projecting neurons significantly reduced food intake at 4 hours in all food intake experiments (n=9-11 for control, n=7-9 for mPFC-LH hM3Dq, two-way ANOVA with Sidak’s multiple comparison test; see supplementary table 2 for details). **G** There was a significant main effect of group on food intake in the light phase (control>mPFC-LH hM3Dq, p=0.035) and a significant group-by-time interaction (p=0.0082). Food intake was significantly reduced in the mPFC-LH hM3Dq group compared to control at 4 hours during the light phase experiment (p=0.0019), at 4 hours during the dark phase experiment (p=0.0461) **H**, and at 4 hours in during the fasting re-feeding experiment (p=0.0287) **I**. **J** Mice were trained with the FED3 for 7 days prior to the progressive ratio task on fixed ratios 1-5 (n=8 for control; n=7 for mPFC-LH hM3Dq). All mice achieved an average correct response rate (% nose pokes to active side of FED3) of greater than 75% and were included in the progressive ratio task. **K** Schematic illustrating the experimental timeline for the progressive ratio task. All mice were tested in an *ad lib* fed state and following a short fast in sessions separated by one week. Mice were counterbalanced to the order of treatment. See methods for details. **L** At 5 hours from the start of the progressive ratio task, the mPFC-LH hM3Dq group reached a significantly lower breakpoint ratio compared to controls in the fasted state (two-way ANOVA with Sidak’s multiple comparison test, p=0.0004) and there was a significant group-by-metabolic state interaction (two-way ANOVA, p=0.0001). **M** Breakpoint ratio did not differ between groups in the fed state (two-way ANOVA, p=0.4893) however there was a significant group-by-time interaction (two-way ANOVA, p<0.0001). **N** In the fasted state there was a main effect of group on breakpoint ratio reached (two-way ANOVA, control>mPFC-LH hM3Dq, p=0.0388) and a significant group-by-time interaction (two-way ANOVA, p<0.0001). Breakpoint ratio was significantly reduced in the mPFC-LH hM3Dq group compared to control from 4-hours to 5-hour time points (two-way ANOVA with Sidak’s multiple comparison test, p<0.05). All data are mean ± SEM. Dots represent individual animals. For statistical analysis, a indicates a significant main effect of group, b indicates a significant interaction, *p<0.05, **p<0.01, and ***p<0.001 for between group comparisons. CNO, clozapine-N-Oxide; FED3, feeding experimentation device version 3; LH, lateral hypothalamus; mPFC, medial pre-frontal cortex.

To assess whether the mPFC-LH pathway suppresses motivated behavior to obtain a food reward (20mg sucrose pellet), we used a FED3 home cage operant conditioning system (22). Mice were trained on FR1-FR3-FR5 schedules until all mice correctly selected the active nose poke 75% of total pokes performed (Fig 4J), and then were subjected to a progressive ratio task following CNO injection (Fig 4K) either in a mildly fasted or ad libitum fed state at the start of the dark phase. DREADD activation of the mPFC-LH circuit suppressed operant responding, as assessed by the breakpoint reached, however the suppression of operant responding only occurred in the acute 3 hr fasted state (Fig 4L-M), when mice ordinarily express increase motivation to obtain a sucrose reward. As expected, fasted control animals reached a higher breakpoint than control animals in the fed state, however, this was not observed after DREADD activation of the mPFC-LH circuit (Fig 4N). Moreover, activation of the mPFC-LH pathway did not affect anxiety-like behavior or locomotion assessed using the elevated zero maze, open field, or light-dark box (Fig S5B-L). There were also no differences in expression of walking, stationary, and rearing behaviors in the open field after DREADD activation. Together with the free feeding experiments described above, our results suggest that activation of the mPFC-LH circuit directly suppresses both appetitive and consummatory feeding behaviors without affecting anxiety-like behavior.

To assess the identity of the mPFC-LH projecting neurons, we injected retrograde AAVs in the LH to express either cre-dependent GCaMP6f or hChR2 in the mPFC-LH pathway in vesicular glutamate transporter-1 cre mice (Vglut1-cre; mPFC^Vglut1^-LH (31)). GCaMP6f expression was observed in mPFC (Fig 5B) and both hand stressor and air puff stress acutely increased activity in the mPFC^Vglut1^-LH pathway (Fig 5C-I). Moreover, exposure to chow, familiar palatable or novel palatable food, but not a wood dowel, produced a significant reduction in mPFCVglut1-LH activity (Fig 5J-K).

**Figure 5.**
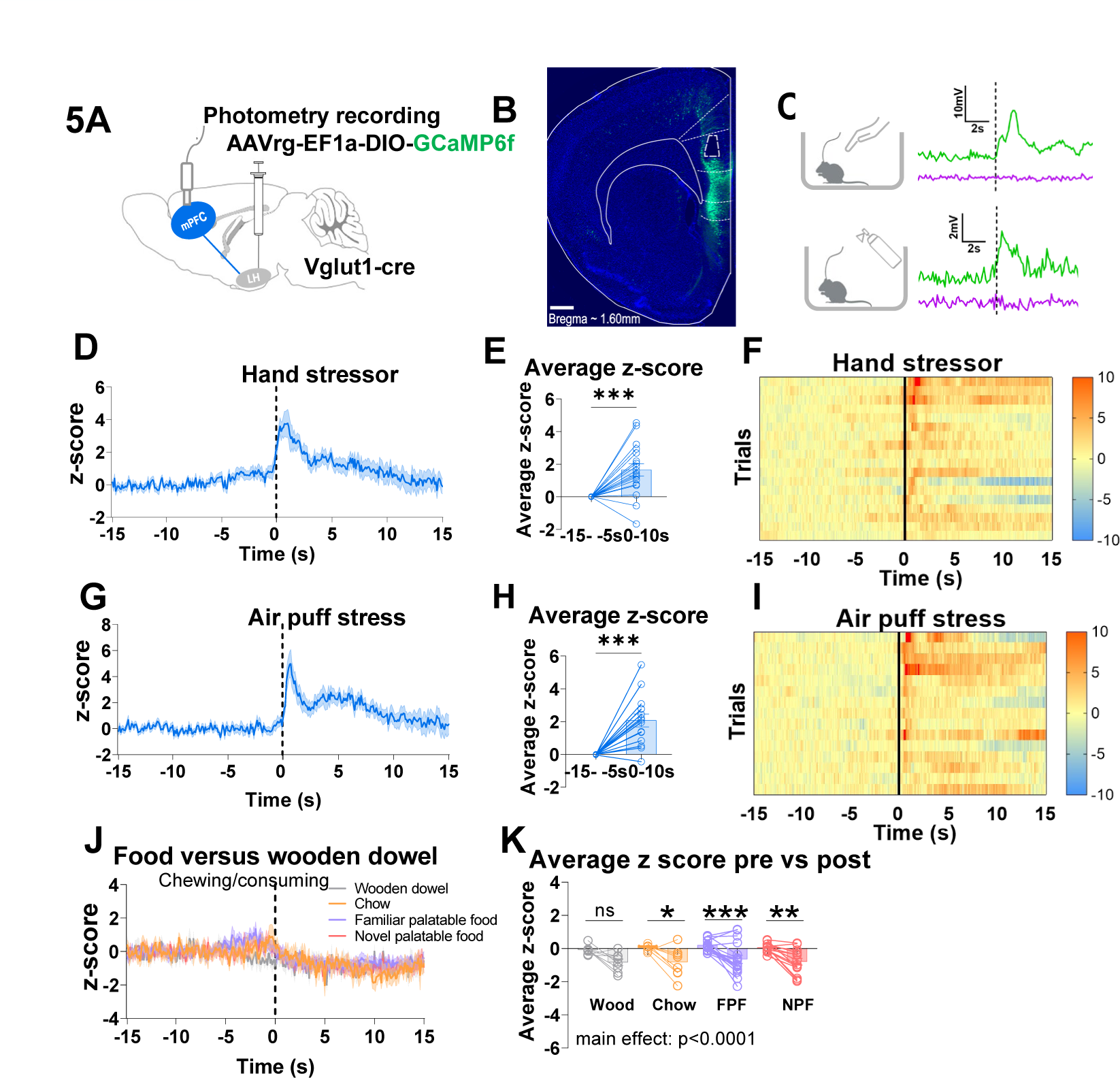
mPFC^Vglut1^-LH projecting neurons respond to acute stressors and are suppressed during food consumption. **A** Schematic illustrating the viral approach used to record from the vesicular glutamate transporter-1 (Vglut1) neurons of the mPFC that project to the LH (mPFC^Vglut1^-LH). Vglut1-cre mice were injected bilaterally into the LH with a retrograde virus encoding for GCaMP6f. **B** Representative image showing GCaMP6f expression in mPFC^Vglut1^-LH projecting neurons with fiber optic cannula placement shown as dashed line. Scale bar is 500µm. **C** Short trace demonstrating calcium-dependent fluorescence associated with GcaMP6f (465 nm) and calcium-independent fluorescence associated with the isosbestic control (405 nm). **D** mPFC^Vglut1^-LH projecting neurons are acutely activated by a hand in cage stressor (n=4 mice, n=3-5 trials per animal, 18 trials in total) **E** Average z-score was significantly greater 0-10 seconds after hand entered cage compared to -15- -5 seconds (baseline period) prior to hand in cage (two-tailed paired t-test, p=0.0003). **F** Heat maps showing individual trial responses to hand in cage test. **G** mPFC^Vglut1^-LH projecting neurons are acutely activated by an air puff stressor (n=5 mice, n=2-4 trials per animal, 15 trials in total). **H** Average z-score was significantly greater 0-10 seconds after air puff delivery compared to -15- -5 seconds (baseline period) prior to air puff (two-tailed paired t-test, p=0.0001). **I** Heat maps showing individual trial responses to air puff stressor. **J** mPFC-LH projecting neuron activity during chewing and consumption of wooden dowel (n=4 mice, n=2 trials per animal,8 trials in total, grey); chow (n=4 mice, n=2 trials per animal, 8 trials in total, orange); familiar palatable food (FPF) (n=4 mice, n=3-5 trials per animal, 17 trials in total, purple); and novel palatable food (NPF) (n=4mice, n=3-5 trials per animal, 14 trials in total, pink). **K** Average z-score 0-15s post chewing wooden dowel did not differ from -15-0s pre chewing. Average z-score 0-15s post chewing and consumption of chow, familiar palatable food and novel palatable food was significantly lower than -15-0s pre consumption (ordinary one-way ANOVA with planned comparisons and Sidak’s multiple comparison test: chow -15-0s>0-15s p=0.0337; familiar palatable food -15-0s>0-15s p=0.0011; novel palatable food -15-0s>0-15s p=0.0037). All data are mean +/- SEM. Circles represent individual trials. Dashed lines at 0 (**D, G** and **J**) and solid lines at 0 (**F** and **I**) indicate time at which event occurred. *p<0.05, **p<0.01, ***p<0.001, ****p<0.0001. LH, lateral hypothalamus; mPFC, medial pre-frontal cortex; ns, not significant; Vglut1, vesicular glutamate transporter-1.

To test the function of the mPFC^Vglut1^-LH circuit with faster temporal dynamics, we employed a wireless optogenetics approach (32), injecting a retro-AAV encoding for channelrhodopsin (ChR2) in the LH and placing a LED probe in the mPFC to photostimulate the mPFC^Vglut1^-LH pathway (Fig 6A, S6A). We confirmed the expression of retrograde hChR2(H134R) throughout the mPFC (Fig 6B). Photostimulation of the mPFC^Vglut1^-LH pathway suppressed feeding after a 14-hour fast when stimulation was paired with returning food to the home cage (Fig 6C, S6B). Next, we photostimulated the mPFC^Vglut1^-LH circuit during a real-time place preference task, to assess the inherent rewarding or aversive nature of this circuit (Fig 6D). Mice spent less time in the side of the arena paired with photostimulation in mPFC^Vglut1^-LH photostimulation group, compared to a light-off control period (Fig 6E), which was not due to differences in locomotor activity (Fig S6E) This suggests that photostimulation of the mPFC^Vglut1^-LH circuit conveys aversive-state information, which may contribute to the reduction in food intake. Based on these results, we hypothesized that photostimulation of the mPFC^Vglut1^-LH circuit would suppress motivation to obtain a sucrose pellet, consistent with our DREADD activation studies. We trained mice in a procedure similar to that described in Fig 4K before testing 3-hour fasted mice with a progressive ratio task under conditions of photostimulation and no stimulation (Fig S6C). Photostimulation of the mPFC-LH^Vglut1^ circuit suppressed the breakpoint reached and active nose pokes (Fig 6F) consistent with results from the DREADDs studies. Together our results suggest the mPFC^Vglut1^-LH circuit responds to acute stress and reduces feeding and motivated reward-seeking behavior by transmitting an aversive signal.

**Figure 6.**
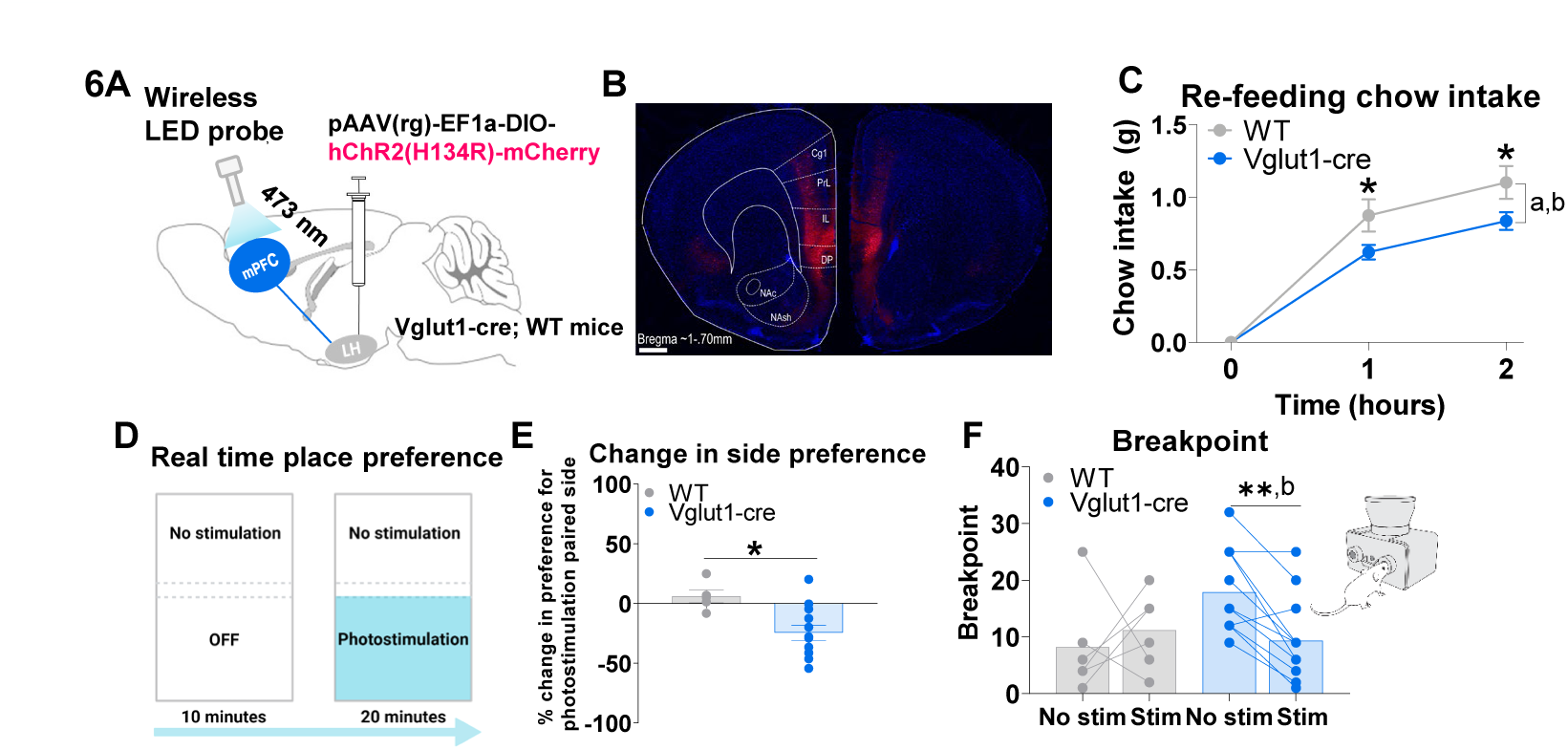
Wireless optogenetic activation of mPFC^Vglut1^-LH projecting neurons suppresses feeding and promotes avoidance. **A** Schematic showing viral strategy and probe used for optogenetic activation of the vesicular glutamate transporter-1 (Vglut1) neurons of the mPFC that project to the LH (mPFC^Vglut1^-LH). Vglut1-cre mice and wild-type (WT) littermates were injected bilaterally into the LH with a retrograde virus encoding for the excitatory opsin hChR2(H134R)-mCherry. A wireless probe with a light emitting diode was inserted into the mPFC. **B** Representative coronal section showing expression of hChR2(H134R)-mCherry (red), with DAPI staining (blue), throughout the mPFC following bilateral injection of retrograde pAAV(rg)-EF1a-DIO-hChR2(H134R)-mCherry into the LH of a Vglut1-cre expressing mouse Cg1, cingulate gyrus 1; DP, dorsal peduncular cortex; IL, infralimbic cortex; NAc, nucleus accumbens core; NAsh, nucleus accumbens shell; PL, prelimbic cortex. Scale bar represents 500µm. **C** Photostimulation of the mPFC^Vglut1^-LH projecting neurons suppresses re-feeding following an overnight fast (n=7 for WT, n=15 for mPFC^Vglut1^-LH, two-way ANOVA with Sidak’s multiple comparison test). There was a significant main effect of photostimulation on chow intake (WT > mPFC^Vglut1^-LH, p=0.0287) and a significant group-by-time interaction (p=0.0118), and significantly reduced chow intake in the mPFC^Vglut1^-LH group compared to WT at 2 hours (p=0.0224) and 4 hours (0.0157). **D** Schematic illustrating the real-time place preference (RTPP) test. Mice were given 10 minutes to explore the RTPP arena before one side of the area was paired with photostimulation for 20 minutes. **E** mPFC^Vglut1^-LH animals reduced their preference for the photostimulation paired side of the RTPP arena (% change from baseline period preference) compared to WT group during the 20-minute RTPP test (n=5 WT, n=12 mPFC^Vglut1^-LH, two-tailed unpaired t-test, p=0.011). **F** Following the progressive ratio task using the FED3, breakpoint ratio was significantly reduced in the mPFC^Vglut1^-LH mice under photostimulation conditions compared to no stimulation (n=6 for WT, n=13 for mPFC^Vglut1^-LH, two-way ANOVA with Sidak’s multiple comparison test, mPFC^Vglut1^-LH No Stim> mPFC^Vglut1^-LH Stim, p=0.0056), and there was a significant group-by-time interaction (p=0.0169). All data are mean +/- SEM. For statistical analysis, a indicates a significant main effect of group, b indicates a significant interaction, *p<0.01 and **p<0.01 for between group comparisons. FED3, feeding experimentation device version 3; LH, lateral hypothalamus; mPFC, medial pre-frontal cortex; Vglut1, vesicular glutamate transporter-1.

## Discussion

The neuronal control of food intake is essential for the maintenance of a healthy body weight and involves a complex interplay between homeostatic, motivational, and cognitive drivers. Here, we used human brain imaging and spectral dynamic causal modelling of effectivity connectivity to predict that the inhibitory influence of a vmPFC to LH pathway is attenuated in a hungry state, compared to sated state. These human studies served as a screening tool to identify a novel neural pathway (mPFC-LH) associated with hunger status and guided our functional preclinical studies. This approach may potentially improve the translation of preclinical studies into impactful clinical outcomes in the long term. In mice, we demonstrated that the mPFC-LH circuit is sensitive to acute stressors and the acute stress associated with novelty suppresses feeding and motivated sucrose seeking.

The mPFC is a brain region implicated in a broad range of responses that includes the encoding of cue- and value-association, and decision making related to reward-seeking (10, 33). Intriguingly, activation of mPFC neurons can either promote or suppress reward responding depending on the projection specific population targeted, including well-studied projections to the nucleus accumbens (NAc), ventral tegmental area (VTA), paraventricular thalamus (PVT) and basolateral amygdala (BLA) (24, 34, 35). Although anatomical mPFC projections to the LH have been previously described (17, 18), the function of this circuit remains to be fully explored, with recent studies suggesting a role in aggressive behavior in response to changing social threats (19) and in impulsive behavior (36). Consistent with previous anatomical reports, we show mPFC neurons project from the PrL, IL and Cg1 subregions of the mPFC directly to the LH. Our results demonstrate that the mPFC-LH circuit, more specifically a mPFC^Vglut1^-LH circuit, is sensitive to stressful stimuli, which produces an inhibitory effect on food intake and motivation to consume sucrose pellets. Specifically, both acute chemogentic activation of the mPFC-LH circuit and optogenetic activation of mPFC^Vglut1^-LH neurons suppresses food intake as well as the motivation to obtain sucrose pellets. We acknowledge that mPFC-LH projecting neurons may send collateral projections to other downstream targets. However, previous studies suggest that mPFC to BLA projections increase feeding (34) and mPFC to NAc projections act to promote reward seeking (35). Our findings are in line with evidence that cortical inputs from the anterior insula cortex to the LH alter feeding behavior and body weight maintenance (26) providing evidence for a unique role of cortical to LH circuits.

The mPFC-LH pathway was engaged by acute stressors in this study, a finding that is supported by a known role of the mPFC in integration (20, 37–40) of acute stress (41–43). Our study, however, moves beyond the broad association between acute stress and mPFC activity to illustrate that acute stress specifically recruits the mPFC-LH pathway to suppress food intake and motivation. Moreover, the activity of this pathway may *predict* acute stress, as shown by the recruitment of mPFC-LH neurons immediately prior to entry into a novel environment or on approach of a novel object, consistent with the notion that mPFC neurons are important in decision-making (44–46) and event or threat prediction (20, 37–40). We acknowledge that this idea requires further evidence and that other factors, such as learning and prediction error, may underlie this effect. Nevertheless, this prediction-like activity diminished over subsequent events, suggesting that prior experience updates the function of this circuit. Similar habituation in response to stress experience has been observed in CRH neurons (47). Our results suggest the mPFC-LH circuit is specifically relevant to acute stress, rather than generalized anxiety, because chemogenetic circuit manipulation had no affect anxiety-like behavior across multiple well-established assays. Further evidence to support this comes from the reduced time spent rearing in the open field after mPFC-LH inhibition, since rearing behavior is normally expressed within a novel environment to survey for potential danger or threat (30). Food consumption, but not food presentation, approach, or wooden dowel chewing reduced mPFC-LH activity and surprisingly, the magnitude of the fall was not related to metabolic state, palatability or food consumed, as has been observed with hunger-sensing AgRP neurons (48, 49). Altered mPFC activity has been observed in response to palatable food consumption in models of binge eating (50), suggesting that this circuit may play an important role regulating emotional state in the absence of homeostatic need. In support of this, mPFC-LH DREADD inhibition increased acute motivated reward-seeking, food approach, and consumption only under conditions of mild stress, such as in a novel environment or in response to a novel food (51) and not during home cage feeding.

There is conflicting data on the role of the mPFC in cue-dependent learning, value association and motivation to seek reward. For example, mPFC activity is associated with the inhibition of motivated, natural, and drug-related reward seeking (24, 52–55). However, opposing results suggest that mPFC activity drives responding for natural and drug-related rewards and reinstatement of cue- or context-induced reward seeking following extinction (34, 35, 56, 57). It is worth reiterating that human neuroimaging studies also report discrepancies in mPFC activity with hyperactivation in both obesity (10, 33) and anorexia nervosa (11, 58). These differences may be largely explained by the projection targets of the mPFC neurons in question, in which corticostriatal pathways increase reward seeking and corticothalamic pathways inhibit reward seeking (35). Our results build upon this model by including cortico-hypothalamic pathways as inhibitory pathways that are specific for food-related reward seeking. Supporting this idea and relevant to the feeding experiments herein, photostimulation of mPFC dopamine 1 receptor-expressing neurons, which project to both striatal and limbic regions, promotes food intake and responding for food rewards (34). Other studies have demonstrated that activation of corticolimbic circuits promotes reward seeking (59). Conceivably, mPFC limbic and striatal projections have similar roles in promoting reward seeking, while thalamic and hypothalamic projections generally act to suppress food intake and motivated behaviors.

These studies raise an important question; does the mPFC target a specific subpopulation of LH neurons to inhibit feeding and motivation? Photostimulation of LH*^Vgat^* neurons, which are molecularly distinct from orexin and melanin concentrating hormone (MCH) expressing neurons, promotes feeding in sated animals, while photoinhibition has the opposite effect (15). In contrast, photostimulation of LH*^Vglut2^* neurons suppresses food intake in fasted animals and is aversive (60). Our chemogenetic and optogenetic approaches directly phenocopy studies manipulating LH*^Vglut2^* neurons. An important recent study shows that LH*^Vgat^*and LH*^Vglut2^* neurons are not sensitive to interoceptive signals of homeostatic state (61), supporting our observations that mPFC-LH neural activity in response to food is not sensitive to metabolic state. We predict that LH*^Vglut2^* neurons are a primary downstream target of excitatory mPFC-LH projecting neurons. Indeed, a recent set of studies show that LH*^Vglut2^* neurons form part of a neural circuit that can suppress hunger-sensing AgRP neurons via a GABAergic population in the dorsomedial hypothalamic nucleus (62). In addition, neurons from the mPFC (PrL and IL) have monosynaptic inputs with LH orexin and MCH neurons, some of which have been shown to express *Vglut2* and are differentially affected by air puff stress, immobilization or novel object presentation and investigation (63).

Given the sensitivity to acute stressors, the mPFC-LH circuit may play an important and underappreciated role in the pathogenesis of psychometabolic conditions. For example, increased mPFC activity is associated with anorexia nervosa (11), excessive dietary self-control (7), and chronic inhibition of specific mPFC projections reduces anorectic symptoms in a rat model of activity-based anorexia (64). On the other hand, decreased activity upon consumption removes this restraint and may, in some circumstances, promote overeating or binge eating. While this might be an important new target to combat overeating associated with the progression to obesity, it’s important to note that human neural network modelling did not identify BMI as a factor affecting mPFC-LH activity.

In summary, we have used neural network modelling from human imaging data, together with neural circuit control in mice, to identify a novel anorexigenic mPFC^Vglut1^-LH pathway involved in the regulation of food intake and motivation. Neural recordings using photometry revealed that this pathway responds to acute stressors and predicts potential threat when animals voluntarily enter a novel environment or approach a novel object. Intriguingly, this stress-sensitive pathway is suppressed in response to food consumption but not affected by metabolic state or palatability, which establishes a novel neural pathway that may provide top-down cortical control over food intake and motivation, independent of hunger or energy needs. Future studies are required to elucidate the environmental, developmental, psychological, and physiological cues affecting this system as they might play important roles in the pathogenesis of opposing psychometabolic conditions such as anorexia nervosa or overeating and obesity.

## Acknowledgments

This work was supported by National Health and Medical Research Council project grants APP1126724 & APP2013243 (ZBA and AVG) and a National Health and Medical Research Council research fellowship APP1154974 (ZBA). We would like to thank Antonio (Nino) Benci at the Monash Instrumentation Facility for help with FED3 production and maintenance.

## Disclosure Statement

The Authors have nothing to disclose

**Video 1** – mPFC-LH GCaMP6s response to repeated pick up at 10x normal speed. The video shows raw collected in mV at the photoreceiver prior to df/f calculations.

**Video 2** – mPFC-LH GCaMP6s response to air puff at 10x normal speed. The video shows raw collected in mV at the photoreceiver prior to df/f calculations.

---

**Figure S1.**
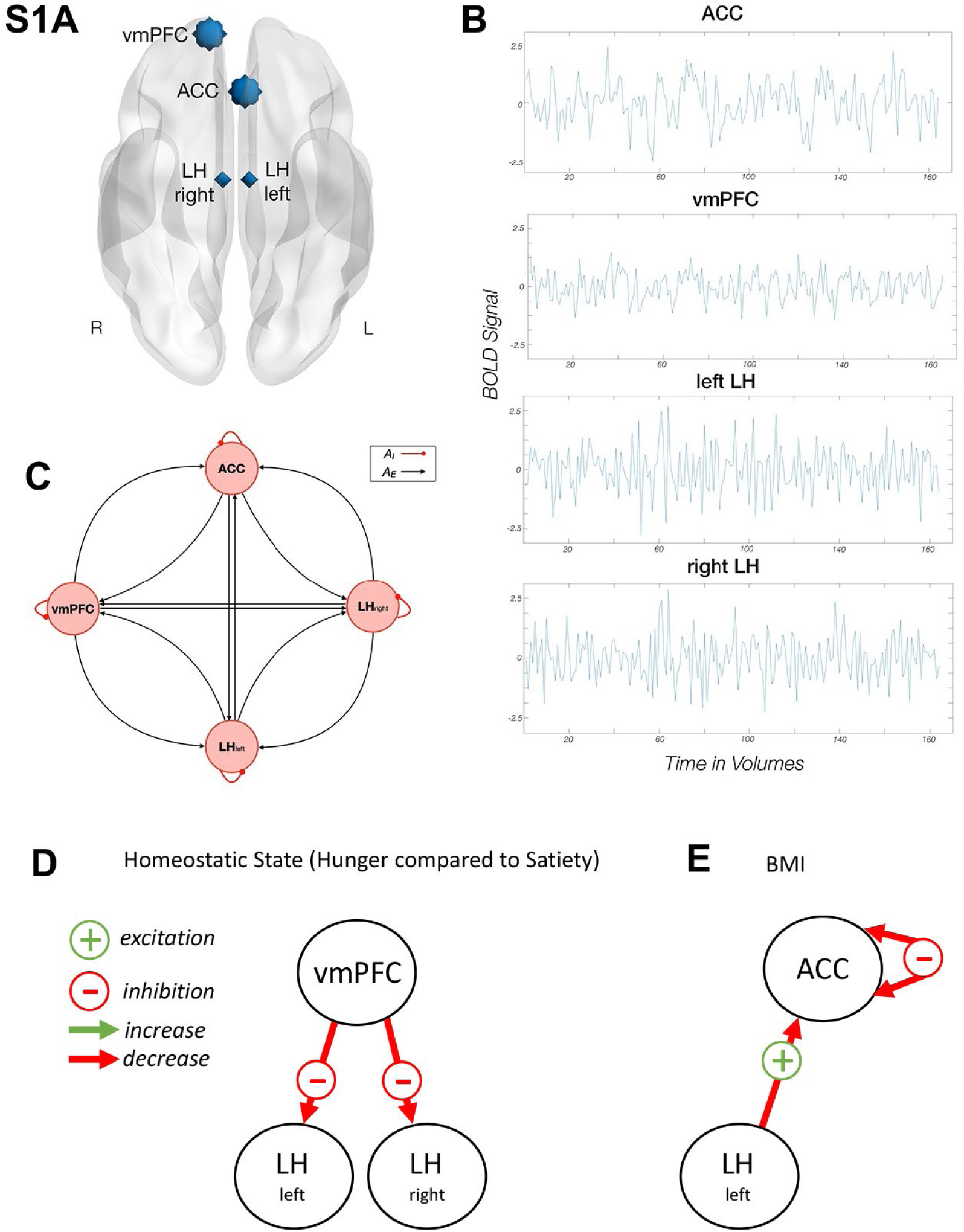
Spectral dynamic causal modeling. **A** Regions of interest (ROIs) for neural network modelling were selected based on data from Harding et al. (2018), which established functional connectivity between the PFC and LH. **B** The time series of the ROIs for an exemplar subject. Anterior cingulate cortex, ACC; Ventromedial prefrontal cortex; vmPFC; LH, Lateral hypothalamus. **C** Schematic showing the ROIs used to estimate spDCMs with the fully connected architecture (i.e., 4^2^ = 16 parameter model; Friston et al., 2014). Second level spDCM results. **D** Effective connectivity demonstrates hunger was associated with decreased inhibition from the ventromedial prefrontal cortex to the bilateral lateral hypothalamus. **E** Effective connectivity demonstrates BMI was associated with decreased excitation from the left lateral hypothalamus to the anterior cingulate cortex (ACC) and decreased inhibition within the ACC. + or – signs code the activity state of connectivity: –, inhibitory; +, excitatory. Colored arrows indicate connectivity strength with a decrease shown in red and an increase in green.

**Figure S2.**
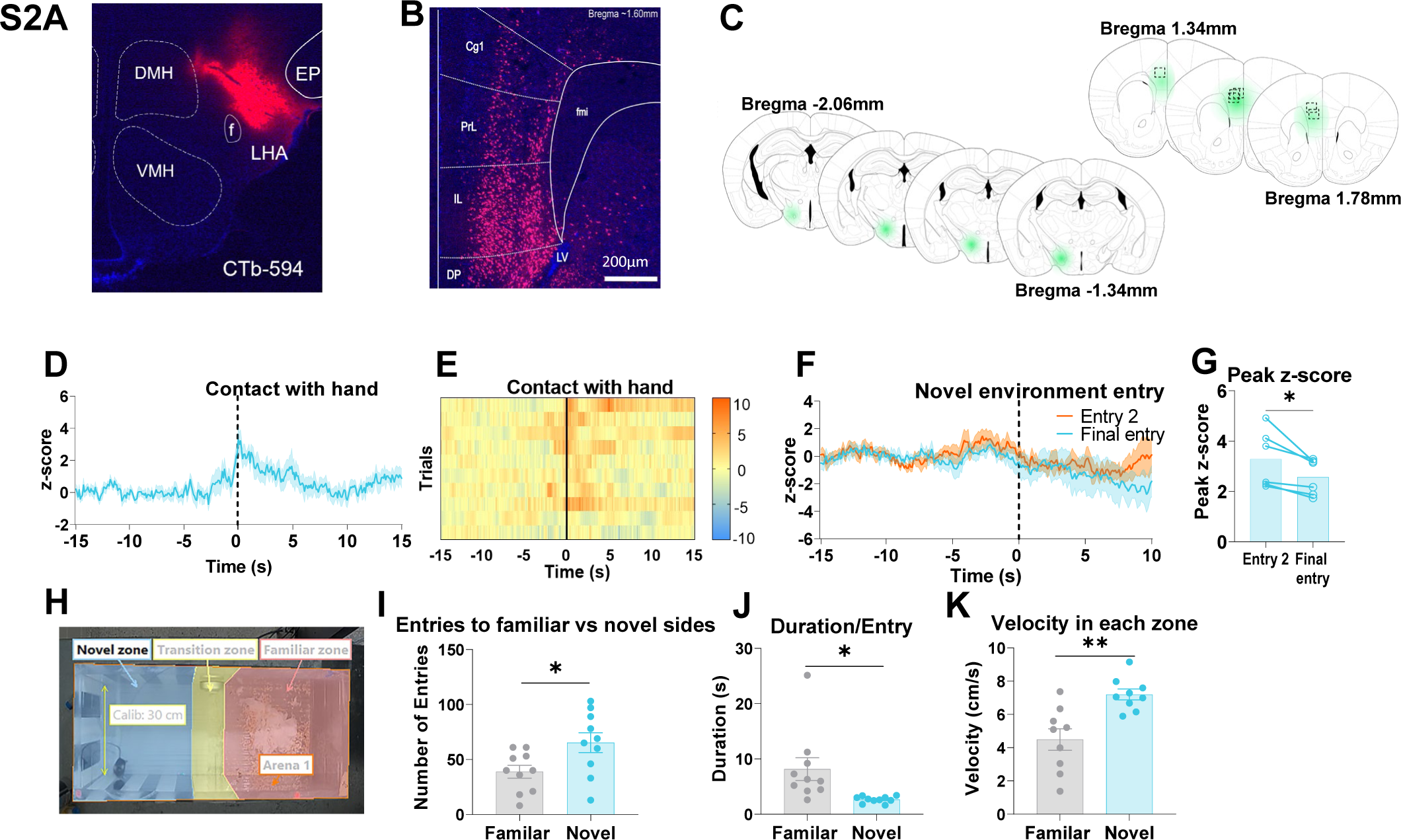
mPFC-LH photometry injection sites and additional behavioral data. **A** Representative coronal section of LH following injection of Cholera-toxin B (CtB) Alexa fluor 594 (red) with DAPI staining (blue). DMH, Dorsal medial hypothalamus; VMH, Ventral medial hypothalamus. **C** Representative coronal section showing transport of Alexafluor 594 (red), with DAPI staining (blue), throughout the Cg1, PrL and IL regions of the mPFC following bilateral injection of CtB-594 into the LH. **C** Schematic illustrating spread of retrograde AAV-cre injected into the LH (and spread of GCaMP7s injected into the mPFC and cannula placement (n=6, darker green indicates overlapping injection sites between animals, dashed boxes indicate cannula position) **D** A subset of mice from the hand in cage experiment in figure 1H-K were picked up by the tail (n= 3 mice, 3-4 trials per animal, 10 trials in total). mPFC-LH projecting neuron activity increased prior to contact with the researcher’s hand (time=0). **E** Heatmap showing individual trials of response to contact with hand in order of trial. **F** mPFC-LH projecting neuron activity varied from early entry into a novel environment (entry 2, orange) to final entry (entry 10-20, blue) (n=6 mice, data from figure 1P-S). **G** Peak z-score was significantly lower on the final entry to the novel environment compared to entry 2 (two-tailed paired t-test, p=0.0252). **H** Screen capture illustrating the 3 zones used to analyze behavior in the novel environment experiment from figure 1P-S (n=9-10 mice, 10-minute trials). **I** Number of entries into the novel zone in the novel environment experiment was significantly greater than entries into the familiar zone (two-tailed t-test, p=0.0244). **J** Duration spent in the novel side during each entry was significantly reduced compared to the duration spent in the familiar side per entry (two-tailed t-test, p=0.0153). **K** Velocity in the novel zone was significantly higher than velocity in the familiar zone (two-tailed t-test, p=0.0018). All data are mean +/- SEM. Circles represent individual trials while dots represent individual animals. Dashed lines at 0 (**D** and **F**) and solid lines at 0 (**E**) indicate time at which event occurred. *p<0.05, *p<0.01, and ***p<0.001.

**Figure S3.**
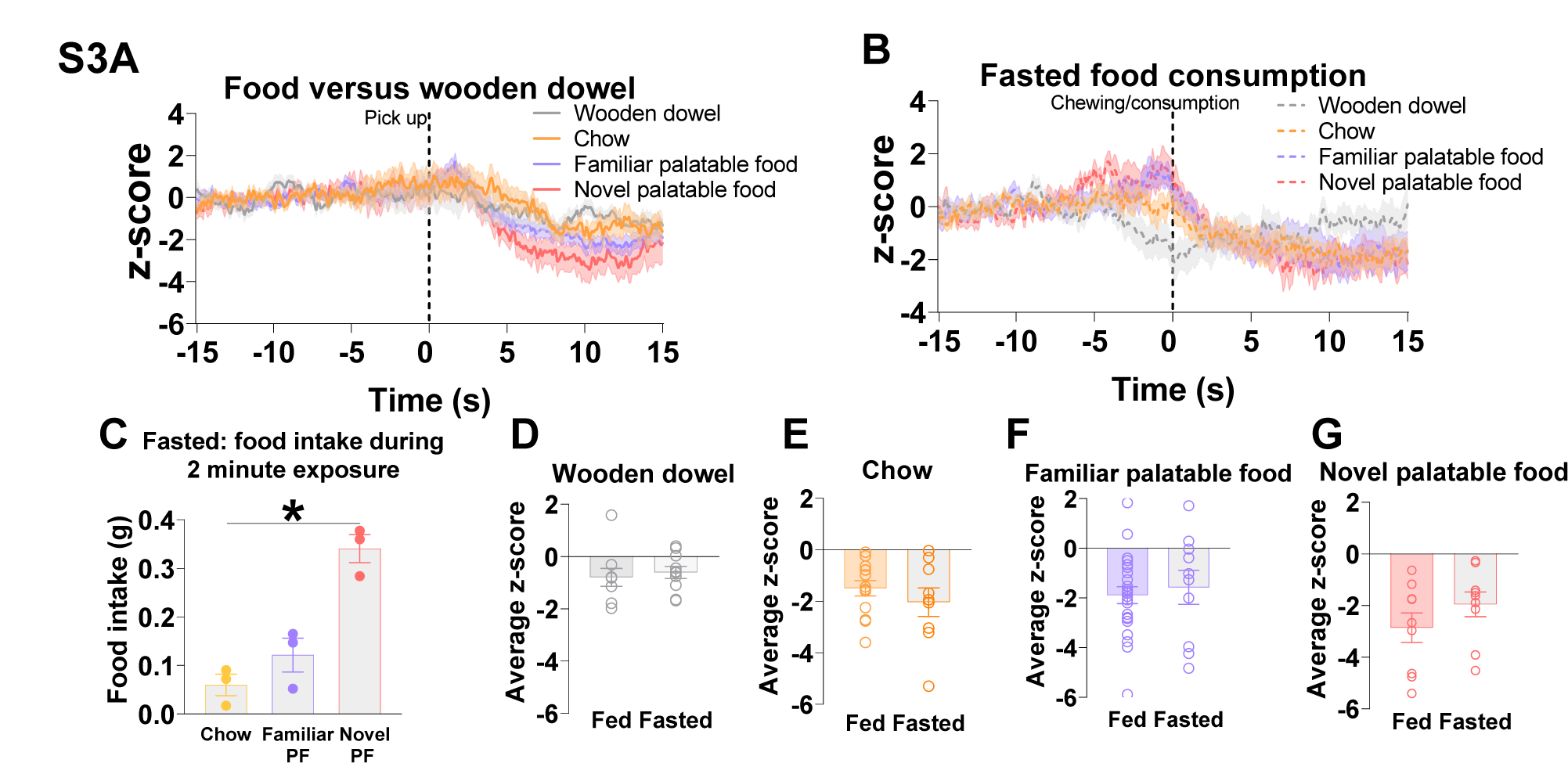
mPFC-LH circuit activity under fed and fasted conditions. **A** mPFC-LH projection neuron activity following pick up of either chow, familiar palatable food, novel palatable food or wooden dowel from figure 2A. **B** mPFC-LH projecting neuron activity following chewing and consumption of wooden dowel (n=3 mice, 2-5 trials per animal, 10 trials total, grey dashed line), chow (n=3 mice, 3-9 trials per animal, 16 trials total, orange dashed line), familiar palatable food (n=3 mice, 2-5 trials per animal, 11 trials total, purple dashed line), and novel palatable food (n=3 mice, 1-4 trials per animal, 9 trials total, pink dashed line) following a short fast. **C** Food intake of chow (orange), familiar palatable food (FPF, purple) and novel palatable food (NPF, pink) over a 2-minute exposure period following a short-term fast (n=3 mice, average over 1-9 trials). Food intake of novel palatable food was significantly greater compared to chow (one-way ANOVA with Tukey’s multiple comparison test, novel palatable food>chow, p=0.0447). **D-G** Average z-score at 5-10s post chewing and consumption was not significantly different in a fed or fasted state. All data are mean +/- SEM. Dashed lines at 0 (**A** and **B**) time at which event occurred. *p<0.05. LH, lateral hypothalamus; mPFC, medial pre-frontal cortex.

**Figure S4.**
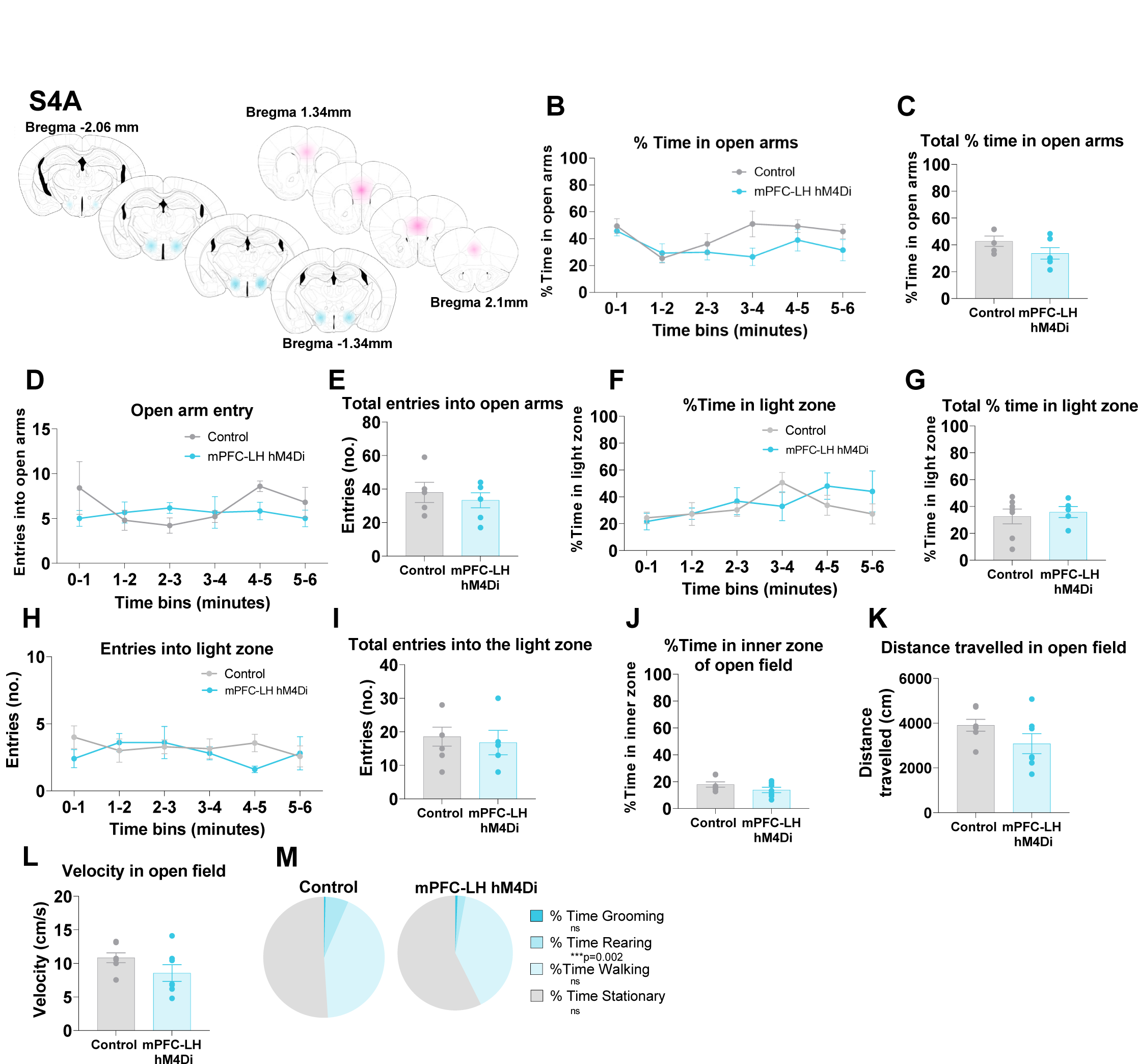
Injection sites and behavioral data for mPFC-LH hM4Di mice. **A** Schematic illustrating spread of retrograde AAV-cre injected into the LH and spread of hM4Di-mCherry DREADDs injected into the mPFC (n=7, darker color indicates overlapping injection sites between animals). **B-M** Mice were placed in the elevated zero maze, light-dark box and open field 30 minutes after intraperitoneal injection (i.p.) of CNO (3mg/kg) (n=5-7 for control, n=6-7 for mPFC-LH hMD4i). mPFC-LH circuit inhibition had no effect on: **B-E** time in the open arms of the elevated zero maze, entries into the open arms; **F-I** time in the light side of the light-dark box or entries into the light zone; or **J-L** time spent in the inner zone, distance travelled or velocity in the open field. **M** mPFC-LH inhibition significantly reduced time spent rearing in the open field (two-tailed t-test, p=0.0002). All data are mean +/-SEM. LH, lateral hypothalamus; mPFC, medial prefrontal cortex.

**Figure S5.**
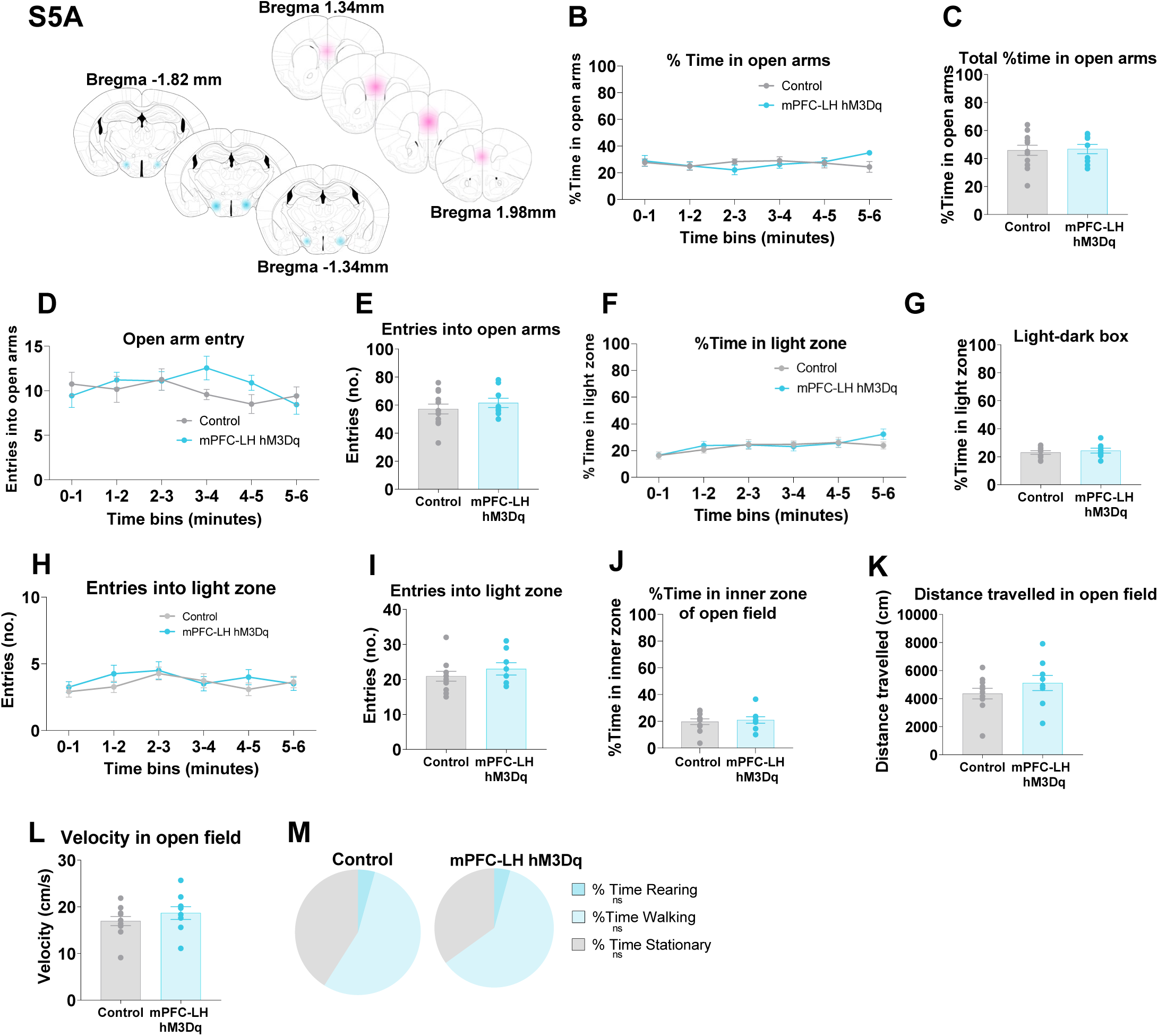
Injection sites and behavioral data for mPFC-LH hM3Dq mice. **A** Schematic illustrating spread of hM3Dq-mCherry DREADDs injected into the mPFC (n=9, darker pink indicates overlapping injection sites between animals). **C-H** Mice were placed in the elevated zero maze, light-dark box and open field 30 minutes after intraperitoneal injection (i.p.) of CNO (1mg/kg) (n=11-12 for control, n=9 for mPFC-LH hM3Dq). mPFC-LH circuit activation had no effect on: **B-E** time in the open arms of the elevated zero maze or entries into the open arms; **F-I** time in the light side of the light-dark box or entries into the light zone; or **G-L** time spent in the inner zone, distance travelled or velocity in the open field; and **M** time spent rearing, walking or stationary in the open field. All data are mean +/- SEM. CNO, clozapine-N-Oxide; LH, lateral hypothalamus; mPFC, medial prefrontal cortex.

**Figure S6.**
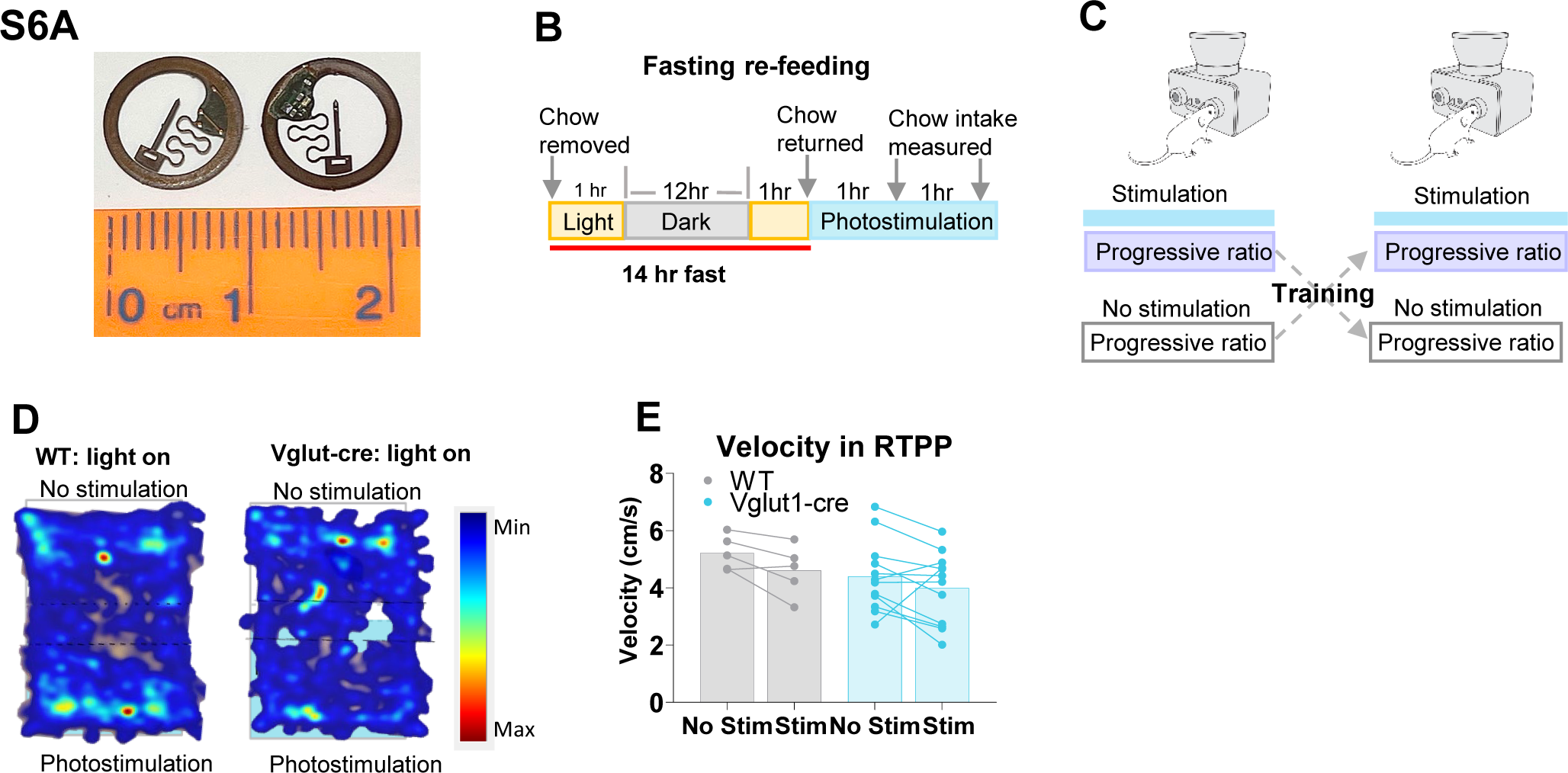
Wireless optogenetic activation of mPFC^Vglut1^-LH. **A** Photograph illustrating wireless antenna with LED probe used in the wireless optogenetics experiments in figure 6. **B** Schematic illustrating experimental timeline for fasting re-feeding with photostimulation. **C** Schematic illustrating the progressive ratio task using FED3 during photostimulation and no stimulation. Mice were counterbalanced to the order of treatment and trained on FED3 between progressive ratio test days. See methods for further details. **D** Representative heat maps from WT (left) and mPFC^Vglut1^-LH (right) animals during the 20-minute RTPP test. **H** Velocity in the RTPP arena did not change significantly between baseline (no-stim) to the 20-minute RTPP test (stim). All data are mean +/- SEM. LH, lateral hypothalamus; mPFC, medial prefrontal cortex; Vglut1, vesicular glutamate transporter-1.

