## Supplemental Table 1 for "Identification of a stress-sensitive anorexigenic neurocircuit from medial prefrontal cortex to lateral hypothalamus"

**Table S1. Summary of statistics; human studies**

| Figure | Statisitical test | n | Model | Effect Size | Posterior probability (>.95) |
|---|---|---|---|---|---|
| **1** | Parametrical Empirical Bayes on estimated single-subject spectral dynamic causal models | n = 53 (fasted), n = 53 (sated) (within-subject) | Hunger Main Effect | decreased inhibition from vmPFC to left LH (-0.101 Hz, 95% CI [-0.01,-0.20]) | **0.96** |
| | | | | decreased inhibition from vmPFC to right LH (-0.11 Hz, 95% CI [-0.02, -0.20]) | **0.98** |
| | | | BMI Main Effect | decreased excitation from the left LH to the ACC (-0.02 Hz, 95% CI [-0.002, -0.04]) | **0.97** |
| | | | | decreased self-inhibition of the ACC (-0.024 Hz, 95% CI [-0.01, -0.04] | **0.99** |
| | | | BMIxHunger Interaction Effect | no effects | **< 0.95** |
