## Supplemental Methods and Materials for "Identification of a stress-sensitive anorexigenic neurocircuit from medial prefrontal cortex to lateral hypothalamus"

**MRI data acquisition**

Resting-state fMRI data were acquired using a 3-Tesla Siemens Skyra MRI scanner equipped with a 32-channel head coil at the Monash Biomedical Imaging Research Centre (Melbourne, Victoria, Australia). For 40 participants, 189 gradient-echo planar images comprising of 44 interleaved, continuous axial slices were collected (repetition time = 2500 ms; echo time = 30 ms; flip angle = 90°; 3 mm isotropic voxels; field of view = 192 mm) during an 8-minute resting-state scan scan. Resting-state sequence changed for the remaining 13 participants: During a total acquisition time of 7.8 minutes, 600 volumes are acquired for each participant and hunger condition using a multiband gradient echo pulse sequence (45 axial slices; time of repetition, TR = 780 ms; echo time, TE = 21 ms, resolution 3x3x3mm). A whole-brain T1-weighted magnetization-prepared rapid gradient-echo structural image was also acquired for each session and each participant (192 sagittal slices; 1 mm isotropic voxels; repletion time = 2300 ms; field of view = 256 mm). Participants were instructed to rest while fixating on a central cross presented on a screen.

**Resting-state fMRI data analyses**

***Preprocessing.*** Functional images were preprocessed using SPM12 (revision 12.2, www.fil.ion.ucl.ac.uk). The preprocessing steps consisted of slice time correction, realignment, spatial segmentation and normalization to the standard EPI template of the Montreal Neurological Institute (MNI), and spatial smoothing using a Gaussian kernel of 6-mm FWHM. No (band-pass) filtering was used except a low-pass filter (of 1/128) that filters the ultra-low frequency scanner drifts^1^. None of the participants exceeded excessive head motion of larger than 3mm.

***ROI selection and time series extraction.*** The identified neural circuit comprised of the ventromedial prefrontal cortex (vmPFC), anterior cingulate cortex (ACC), and bilateral lateral hypothalamus (LH). Following the resting-state fMRI scan of each session, participants completed a task-based fMRI session, during which they were required to make realistic choices between unhealthy and healthy drinks. The vmPFC and ACC ROIs were functionally defined based on results during this task-based fMRI session (Table 1)^2^. We defined the bilateral LH ROIs according to Baroncini et al. (2012)^3^ (Montreal Neurological Institute coordinates; x = ± 6, y = -9, z = -10) using 2-mm-radius spheres (Table 1). The seeds were as such spatially separated by > 6mm (i.e., > 1mm after smoothing).

**Table 1.** ROIs used for spDCM analyses

| Region | MNI Coordinates | | | Sphere size | Source |
| --- | --- | --- | --- | --- | --- |
|  | x | y | z |  |  |
| vmPFC | 12 | 58 | 22 | 6mm | Harding et al. (2018) |
| ACC | -4 | 32 | 30 | 6mm | Harding et al. (2018) |
| Left LH | -6 | -9 | -10 | 2mm | Baroncini et al. (2012) |
| Right LH | -6 | -9 | -10 | 2mm | Baroncini et al. (2012) |

Abbreviations: ACC, anterior cingulate cortex; vmPFC, ventro-medial prefrontal cortex; LH, lateral hypothalamus; MNI, Montreal Neurological Institute.

To extract BOLD fMRI time series corresponding to the aforementioned ROIs, the pre-processed data was used to establish the residuals of a General Linear Model (GLM). Six head motion parameters and WM/CSF signals were added to the GLM as nuisance regressors. Finally, we selected the MNI coordinates as the center of a 6-mm sphere to compute the subject-specific principal eigenvariate and correct for confounds.

**Spectral Dynamic Causal Modeling**

The spDCM analyses were performed using the functions of DCM12 (revision 7196) implemented in SPM12. In order to address our main hypotheses, we focused on spDCM analyses that assessed (1) changes in effective connectivity of fasting versus satiety condition independent of BMI (main effect of fasting), (2) changes in effective connectivity modulated by BMI (main effect of BMI), and (3) changes in fasting-related effective connectivity modulated by BMI (BMI-by-fasting interaction).

***First Level spDCM Analysis***

On the first-level, a fully-connected model was created for each participant and each session (i.e.4^2^ = 16 connectivity parameters, including seven inhibitory self-connections). Next, we inverted (i.e. estimated) the DCMs using spectral DCM, which fits the complex cross-spectral density using a parameterized power-law model of endogenous neural fluctuations^1^. This analysis provides measures of causal interactions between regions, as well as the amplitude and exponent of endogenous neural fluctuations within each region^1^. Model inversion was based on standard variational Laplace procedures^4^. This Bayesian inference method uses Free Energy as a proxy for (log) model evidence, while optimizing the posterior density under Laplace approximation of model parameters.

***Second Level spDCM Analysis***

To characterize how group differences in neural circuitry were modulated by BMI and Hunger condition, hierarchical models over the parameters were specified within a hierarchical Parametric Empirical (PEB) framework for DCM^5^. Three models we used as follows: first, we investigated the group difference between fasted versus sated conditions and in this PEB analysis, we used BMI, age and gender as regressors of no interest. Second, we were interested associating effective connectivity with BMI and in this PEB analysis, we used group factor (fasted versus sated), age and gender as regressors of interest. Lastly, we were interested in interaction between group factor (fasted versus sated) and BMI and in this PEB analysis, we used BMI, group factor (fasted versus sated), age and gender as regressors of no interest.

For each of the presented models, all behavioural regressors were mean-centred so that the intercept of each model was interpretable as the mean connectivity. Bayesian model reduction was used to test all reduced models within each parent PEB model (assuming that a different combination of connections could exist^5^ and ‘pruning’ redundant model parameters; parameters of the best 256 pruned models (in the last Occam’s window) were averaged and weighted by their evidence (i.e. Bayesian Model Averaging) to generate final estimates of connection parameters. To identify important effects (i.e., changes in directed connectivity), we compared models (using log Bayesian model evidence to ensure the optimal balance between model complexity and accuracy) with and without each effect and calculated the posterior probability for each model, as a softmax function of the log Bayes factor. We treat effects (i.e., connection strengths and their changes) with posterior probability >0.95 as significant for reporting purposes.

**Animal Studies**

Experimental conditions

For DREADDs experiments, mice were group housed following surgery in groups of 2-3 for the duration of behavioral testing (elevated zero maze, light-dark box, open field). Mice were then individually housed for all food intake experiments, feeding cage experiments and operant conditioning experiments to allow for accurate measurement of consumption and equal access to food with an acclimation period of at least one week. For optogenetics and fiber photometry experiments, mice were individually housed following surgery and throughout experiments to prevent damage to the implanted probe or cannula. Optogenetics experiments were conducted within the light phase. DREADDs experiments and fiber photometry experiments were conducted through the dark phase with mice on a reverse light cycle. Mice were handled for at least 5 min each day for five consecutive days leading up to behavior experiments.

DREADD experiments

For activation of DREADDs, mice were injected in the intraperitoneal cavity with clozapine-n-oxide (CNO) (hM3Dq DREADDs 1mg/kg, hM4Di DREADDs 3mg/kg; Sigma Aldrich, in saline) 30 minutes prior to each behavioral or feeding experiment. For Fos analysis experiments CNO was delivered 90 minutes prior to euthanizing the animals.

Optogenetic stimulation

For all optogenetic experiments light stimulation was delivered at 20Hz, 3 seconds on 1 second off using wireless LED probes (Neurolux; IL, USA) as described under surgical procedures above. An electromagnetic field was constructed around an animal cage (440mm X 340mm with wall height 200mm) and connected to a power distribution control box (12Amps; 10W; Neurolux) which delivered radio frequency power to the wireless LED probes. Prior to each experiment the antenna was tuned to the power distribution control box to ensure the impedance of the power source and antenna matched. Each antenna was tuned to ensure a standing wave ratio of <1.3 and an impedance of 50Ω. Blue light delivery (470nm) was in the range of 1-5mW as calculated using the irradiance calculator provided by Neurolux.

Viral and surgical procedures

Mice were anesthetized using isoflurane (5% for induction, 2% for maintenance) and positioned on a stereotaxic frame (Stoelting, IL, USA). A Neuros syringe (Hamilton, Reno, NV, USA) was used to deliver virus to the target regions; bilateral LH (-1.3mm Bregma; +/- 1.0mm lateral; -4.9mm ventral from surface of brain) and mPFC (+1.55mm and +1.75mm Bregma; +/- 0.3mm lateral; -1.75mm ventral from surface of brain). Following surgery mice were given 3 weeks to recover and to ensure viral expression before experiments commenced. For viral tracing experiments 100nl of pAAVrg-CAG-GFP or pAAVrg-Syn-ChR2(H134R)-GFP was injected bilaterally into the LH. To express either hM3Dq or hM4Di DREADDs in the mPFC to LH projection neurons AAVrg-pgk-cre was injected bilaterally into the LH at a volume of 100nl before the cre-dependent viral construct encoding for the hM3Dq or hM4Di DREADDs or a reporter virus for controls (AAV5/TRUFR-eGFP or pAAV5-hSyn-DIO-mCherry) were injected bilaterally into the mPFC at a volume of 300nl (75nl/site). For optogenetics experiments a retrograde cre-dependent viral construct was injected (AAVrg-EF1a-DIO-hChR2(H134R)-mCherry) bilaterally into the LH of Vglut1-cre mice and WT littermates at a volume of 100nl. A wireless antenna (Fig 5A; 9.8mm diameter, 1.3mm thickness, 6mm probe length; 30 mg weight; Neurolux) was then implanted above the mPFC (+1.65mm Bregma; + 0.25mm or -0.25 lateral; -1.7mm ventral from surface of brain) unilaterally and fixed to the skull of the mouse using the procedure previously described^6^. For fiber photometry experiments in wild type mice AAVrg-pgk-cre was injected bilaterally into the LH at a volume of 100nl and a cre-dependent virus expressing jGCaMP7s (pGP-AAV9-syn-FLEX-jGCaMP7s-WPRE) was injected unilaterally into sites of the mPFC, to ensure full jGCaMP7s coverage from retrograde cre expression (+1.55mm and +1.75mm Bregma; +0.3mm lateral; -1.8mm and -1.6mm ventral from surface of brain), at a volume of 75nl/site. For Vglut1-cre photometry experiments a cre-dependent retrograde virus encoding for GCaMP6f (AAVrg-EF1a-DIO-GCaMP6f-P2A-nls-dTomato) was injected bilaterally into the LH at a volume of 150nl. Following this, a fiber optic cannula (400µm core, NA 0.48, MF1.25 400/430-0.48, Doric Lenses) was implanted above the mPFC (+1.65mm Bregma; + 0.25mm lateral; -1.7mm ventral from surface of brain) and was secured using dental cement (GBond, GC America).

**Table 2.** Viral constructs

| **Virus** | **Producer** | **Source** |
| --- | --- | --- |
| pAAVrg-CAG-GFP | Boyden | Addgene: #37825 |
| AAV5/TRUFR-eGFP |  | UNC Vector Core |
| AAVrg-pgk-cre | Aebischer | Addgene: #24593 |
| pAAV5-hSyn-DIO-hM3D(Gq)-mCherry | Roth | Addgene: #44361 |
| pAAV5-hSyn-DIO-hM4D(Gi)-mCherry | Roth | Addgene: #44362 |
| pAAV5-hSyn-DIO-mCherry | Roth | Addgene: #50459 |
| AAVrg-EF1a-DIO-hChR2(H134R)-mCherry; | Deisseroth | Addgene: #20297 |
| pGP-AAV9-syn-FLEX-jGCaMP7s-WPRE | Kim | Addgene: #104491 |
| AAVrg-EF1a-DIO-GCaMP6f-P2A-nls-dTomato | Ting | Addgene: #51083 |

Immunohistochemistry

After experiments, mice were deeply anesthetized and then transcardially perfused with 0.05M phosphate buffered (PB) saline followed by 4% *w/v* paraformaldehyde (PFA) in 0.1 M PB. Brains were post-fixed in 4% PFA in 0.1 M PB overnight before being transferred to 30% sucrose solutions. Coronal sections of the whole brain were cut at 30 μm and a one in four series of tissue sections was washed in 0.1 M PB three times before being blocked in 1% H_2_O_2_ in 0.1 M PB for 15 min and then washed again. Tissue was then placed in 0.3% Triton X-100 in 0.1 M PB and 4% normal horse serum for one hour before an overnight incubation at 4°C in primary antibody: anti-mCherry (chicken, 1:1000, Abcam) and anti-cfos (rabbit, 1:200, Santa cruz). Tissue sections were then washed again before a 90 min incubation in secondary antibody at room temperature; donkey anti-rabbit IgG AlexaFluor 488 (1:400; Thermo Fischer Scientific) or goat anti-chicken IgG AlexaFluor 594 (1:400; Thermo Fischer Scientific). The sections were then mounted with hard set mounting medium containing 4’, 6-diamidino-2-phenylindole (DAPI) (Vector) used to counterstain DNA. For Fos counting in hM3Dq and hM4Di DREADDs experiments three coronal sections per animal of cfos stained tissue were imaged and the cingulate gyrus 1, prelimbic cortex, and infralimbic cortex regions of the mPFC were analyzed for % co-expression of reporter protein and Fos using ImageJ.

Behavioral testing

All behavior experiments were video recorded and behavioral tracking analysis was performed by an experimenter blinded to treatment using Ethovision version 14.0.1322 (Noldus Information Technology; NL). The elevated zero maze, light dark box and open field were used to assess anxiety-like behavior as well as novel environment and novel food stress, as described in the supplemental methods. More detailed behavior, such as sniffing, approaching and eating food (as measured in the novel and familiar palatable food intake experiments) was scored manually by a researcher blinded to treatment group.

*Elevated zero maze*

The elevated zero maze (500mm diameter, 260mm height) has two open arm sections (50mm x 330mm) and two closed arm sections (50mm x 330mm with 130mm high barriers). Mice were placed in the center of an open arm and allowed to explore the apparatus for the duration of the test (6 minutes). The percentage time spent in the open arms and number of entries into the open arms reflects the cumulative time in, and entries into, both open arm sections of the maze. Mice that did not remain on the maze for the full duration of the trial were excluded from the analysis.

*Light dark box*

The light-dark box (285 mm wall height) was constructed from wood and consisted of a larger light chamber (480 mm X 295 mm; painted white) and a smaller dark chamber (150 mm X 295 mm; painted black) separated by a small opening. Mice were placed in the center of the dark chamber and allowed to move freely between the two chambers for the duration of the test (6 minutes). Time in the light chamber and the number of entries into the light chamber were measured.

*Open field*

Mice were placed in the outer ring of a large open field (800 mm diameter with wall height 300 mm) and allowed to freely move about the apparatus for the duration of the test (6 minutes). Total distance travelled and average velocity were measured.

*Familiar palatable food in novel environment*

Mice were conditioned to Froot Loops (Kellogs; Australia) three times in the week leading up to the experiment (a single Froot Loop was placed into a plastic weigh boat in the home cage on three occasions, the plastic weigh boat remained in the cage until test day). On the test day, mice were injected with CNO 30 minutes before being transferred to a novel test room. Three Froot Loops (90mg) were placed into the weigh boat from the home cage. The weigh boat was then placed into a novel environment before mice were placed at the opposite end of the box and given 10 minutes to explore the arena. Weight of food consumed was measured, along with the time spent approaching, sniffing and eating, number of eating bouts, latency to first eating bout, and time spent in the food zone (20mm radius around the Froot Loops).

*Novel palatable food in familiar environment*

Mice were conditioned to an experimental box (200 mm X 350 mm, with wall height 350 mm) for 20 minutes per day per day for 3 days in a test room different to the room used for the familiar food test described above. On the first day of conditioning, the home cage of each mouse was placed inside the experimental box for 20 minutes. On the second day mice were placed inside the experimental box with home cage bedding for 20 minutes and a plastic weigh boat. On the final day of conditioning mice were placed into the experimental box with just the plastic weigh boat for 20 minutes. On the test day, mice were injected with CNO 30 minutes before being transferred to the test room. Three 30-50mg pellets of novel high fat/high sugar rodent chow (23% fat, Specialty Feeds: SF04-001, Western Australia) were placed into the plastic weigh boat and placed at one end of the experimental box, mice were placed at the other end of the box and given 10 minutes to explore the environment.

*Real-time place preference*

Mice were given 10 minutes to explore the real-time place preference apparatus (385mm X 245mm with wall height 170mm) before 20 minutes of stimulation (20Hz, 3 seconds on, 1 second off) was paired with one side of the apparatus. We wired the apparatus so that sufficient power to light the LED would only be delivered to one half of the box (stimulation side). Aluminum foil was used to cancel the electromagnetic field on the no stimulation side of the box, however low levels of light were still observed at the very edge of the stimulation side so the center zone was labelled as a neutral zone and not included in the analysis. Preference for stimulation zone was calculated before and during stimulation based on time spent in each zone.

*Operant conditioning*

Feeding experimental devices 3 (FED3; Open Ephys, Portugal) were used for operant conditioning. FED3s were filled with sucrose pellets as rewards (Test Diet Sucrose tablet 20 mg, Richmond, IN, USA). In both DREADDs and optogenetics experiments mice were trained for 7-10 days beginning at a fixed ratio 1 (FR1) for two hours for three days, then to FR3 for two hours for two days, then to FR5 for two hours for two days or until they reached a correct response rate of 75% or above (Figure 2N). For hM3Dq and hM4Di DREADDs experiments, a progressive ratio task (PR; where nose pokes required to obtain a sucrose pellet increase exponentially) was performed in fed (*ad libitum* with access to chow during the PR session) and fasted (three hour fast) states in the dark phase for 5 hours, separated by one week. Active and inactive pokes, pellets acquired and breakpoint – the number of nose pokes at which responding ceases, were recorded throughout each session. Mice were counterbalanced to the order of treatment (fed or fasted) and in the week between PR sessions mice were trained on two days at FR5 to ensure their response rate was maintained above 75%. The hM4Di DREADDs group also underwent an additional two, two-hour PR sessions, 4 weeks after the first PR sessions in home cage and novel cage. Mice were counterbalanced to the order of treatment (home cage or novel cage) and each PR was separated by one week. Before this, the hM4Di DREADDs cohort were trained on FR5 for three days to ensure their response rate was above 75%. Mice were fed *ad libitum* in the lead up to each PR session and injected with CNO 30 minutes prior to the FED being placed into the cage. For the home cage session, mice remained in their home cage in their holding room for the full two-hour PR task, however chow was removed immediately before the session. For the novel cage PR session, 10 minutes before the PR task, mice were transferred to a novel test room, and into a novel cage which was lined with a novel paper bedding and contained a water sipper but no chow. A clean FED3 was then placed into the cage. Between each PR (home cage or novel), the hM4Di DREADDs group were again trained for two days at FR5 to ensure their response rate continued to remain above 75%. For optogenetics experiments a PR was performed under conditions of no photostimulation and photostimulation, and mice were counterbalanced to order of treatment. Each PR session lasted 5 hours and mice had ad lib access to chow during this period. Between each session, optogenetics mice were maintained for two days at FR5 to ensure their response rate was maintained above 75%.

*Fasting refeeding*

Mice were fasted 1-hour before the onset of the dark phase and food (standard chow) was returned one hour following the start of the next light phase (14 hour fast). Food intake was measured at two hours and 4 hours.

*Light-phase food intake*

Mice were injected with CNO and food (standard chow) was removed from the home cage one hour from the start of the light phase. Food was returned 30 minutes later and food intake was measured at two hours and 4 hours.

*Dark phase food intake*

To minimize variability in *ad libitum* food intake in the lead up to the dark phase, food (standard chow) was removed from the home cage 1 hour before the onset of the dark phase and returned three hours later (three hour fast). Mice were injected with CNO 30 minutes before food was returned. Food intake was measured at two hours and 4 hours.

Fiber photometry recording

Mice were acclimated to the photometry recording room and connected to the tethered fiber optic cables cord (400µm core, 0.48 NA; MF1.25 connector; Doric Lenses) in their home cage within the photometry recording room for 1-hour a day for five days leading up to behavior experiments to minimize stress associated with connecting the probe to the patch cord. Fiber optic cannulas were cleaned with 70% ethanol prior to connecting them to the patch cords to minimize contamination of the signal. Two excitation wavelengths, 465nm (Ca2+-dependent signal) and 405nm (Ca2+- independent isosbestic control signal for movement artefacts) emitted from LEDs (Doric Lenses, Canada) were delivered to a dual-fluorescence minicube (4 port fluorescence minicube, FC connectors; Doric Lenses, Canada) before connecting to a rotary joint (pig-tailed 1x1 Fiber-optic rotary joint, 400µm core, 0.48 NA, FC connector; Doric Lenses, Canada) which allow for uninterrupted movement of the patch cord attached to the fiber optic cannula. GCaMP7s or GCaMP6f and isosbestic fluorescence wavelengths were collected using the same fiber optic cable and filtered back through the dichroic ports of the dual-fluorescence minicube into a femtowatt photoreceiver (model no. 2151; Newport). Synapse software (Tucker-Davis Technologies, FL, USA) controlled and modulated the excitation (465 nm: 210 Hz, 405 nm: 530 Hz), as well as demodulated and low-pass filtered (4 Hz) in real-time via the fiber photometry processer (RZ5P;Tucker-Davis Technologies). LED power was adjusted for each mouse to achieve approximately 200mV and 100mV output for GCaMP and isosbestic control signals, respectively. Synapse software also integrated high-definition video recordings of mouse behavior during experiments to allow behavioral events to be accurately aligned with the fluorescent signals.

Fiber photometry behavior experiments

*Hand in cage*

Mice were connected to the patch cord and left undisturbed for 10 minutes in their home cage before the experiment commenced. The researcher then placed their gloved hand into the cage within view of the mouse for 1-2 seconds. This was repeated three times at intervals of 5 minutes. A subset of mice were picked up by the tail after initial exposure to the hand.

*Air puff*

Mice were connected to the patch cord and left undisturbed for 10 minutes in their home cage to collect baseline recordings, following which they were exposed to an air puff delivered using a can of compressed air. Pressured air at 70psi was directed towards the tail and any errors with direction of air puff delivery were excluded from the data. Air puffs were delivered for 1 second at regular intervals of 5 minutes, three times per session. Air puff delivery was marked by the researcher using a TTL trigger.

*Novel environment*

Mice were conditioned to one side of a two-chambered box (700mm X 250mm with 250mm wall height), for 30 minutes per day, for 5 consecutive days leading up to the test day. The two chambers were separately by a plastic divider, with the familiar chamber being smaller than the novel chamber (300mm X 250mm vs 400mm X 250mm, respectively). During conditioning, mice were placed in the familiar chamber with a handful of their home cage bedding on the floor. In addition, the familiar chamber was paired with a strawberry scent (50ul, strawberry essence) which was pipetted at the base of the outer wall of the familiar chamber prior to every conditioning or test session. The walls of the familiar chamber were visually distinct to the novel chamber (black and white vertical stripes and grey horizontal panels, respectively) and the floor of the novel chamber was covered in plastic and was not paired with a scent. On the test day mice were placed in the familiar chamber, with the novel chamber blocked by the plastic divider for 20 minutes. The divider was then removed and mice had free access to move between the novel and familiar chambers for 20 minutes. Entries into the novel chamber were time stamped using video recordings from the test day. An entry was determined to be when all four paws of the animal had stepped onto novel chamber floor. For analysis of mPFC-LH activity at the beginning and end of the novel environment test session we compared the peak z score from the 2^nd^ and last entry - note the 1^st^ novel environment entry was removed from all analyses to avoid any potential confounding activity related to removing the central divider.

*Presentation of food*

Prior to food presentation experiments, mice were conditioned to a Froot loop (familiar palatable food) in their home cage inside a plastic testing dish 3 times over the week leading up to the initial experiment and then again once, two days before subsequent test days. Novel palatable foods consisted of: Reeces peanut butter chips, Cadbury milk chocolate chips, Cadbury dark chocolate chips, Cadbury white chocolate chips, Cadbury caramel flavored chips, 30mg pieces of pink high fat/high sugar rodent chow (23% fat, Specialty Feeds: SF04-001, Western Australia) and white “western diet” rodent chow (22% fat, Specialty feeds: SF00-219, Western Australia). Novel palatable foods were distinct in shape (chips cut into pieces), color and flavor but of similar caloric density and macronutrient profile to one another. Each mouse received each novel palatable food only once. Mice either had ad libitum access to chow (fed) or had food removed 1-hour before the onset of the dark phase and were tested 2-hours into the dark phase (3-hour fast). On test day mice were given a 10-minute baseline period, and were then habituated to a familiar object and plastic testing dish being placed into the cage. This occurred 5-10 times over a 5-minute period to reduce any response to a hand or object entering the cage. Food (regular chow, familiar palatable food or novel palatable food in a random order) or a piece of wooden dowel was then placed into the plastic testing dish and mice were given 2 minutes to approach and consume the food. In each session mice were exposed to chow and familiar palatable food twice, and to either one or two novel palatable foods or wooden dowel so as to avoid satiation.

Fiber photometry analysis

Following recording, behavioral events were aligned using Open Scope software (Tucker-Davis Technologies) with the researcher blinded to the acquired fluorescent signals. Custom-written python code was used to integrate and analyze the fluorescent signals during behavioral events or researcher-delivered stimuli. Ca2+ and isosbestic signals during recording sessions were extracted and down sampled (10 samples/s). The isosbestic signal was fitted to the Ca2+-dependent signal using a least squares polynomial fit and delta F/F was calculated using the equation: (Ca2+-dependent signal - fitted isosbestic signal)/fitted isosbestic signal to correct for photobleaching of the signal and movement artefacts. Due to the short time frame of each experiment (15-45 minutes) photobleaching of the GCamp7s signal was negligible. Z-score normalization was then calculated from the delta F/F for each trial or behavioral event using the equation: z = (F-Fµ)/Fσ where F is the signal and Fµ and Fσ are the mean and standard deviation of the baseline signal. The baseline was determined to be -15 to -5, or -10 to -5 seconds prior to each behavioral event/stimulus presentation. Average z-scores for each trial were determined and statistically analyzed in Graph Pad Prism, V 9.1.2.
