## Supplemental Table 2 for "Identification of a stress-sensitive anorexigenic neurocircuit from medial prefrontal cortex to lateral hypothalamus"

**Table S2. Summary of statistical tests; animal studies**

| Figure | Statisitical test | n | p-value | Post test | p-value |
|---|---|---|---|---|---|
| **1J** | Two-tailed paired t-test (t=5.648, df=14) | n=5 mice, 3 trials per animal, no. of pairs=15 | **<0.0001** | | |
| **1N** | Two-tailed paired t-test (t=4.503, df=11) | n=4 mice, 3 trials per animal, no. of pairs=12 | **0.0009** | | |
| **1R** | Repeated measures one-way ANOVA | n=6 mice, 9 trials per animal, 54 trials in total | | Dunnet's multiple comparison test | |
| | F (2.328, 123.4)=16.56 | | **<0.0001** | -10- -5s vs. -5-0s | **0.0015** |
| | | | | -10- -5s vs. 0-5s | **0.4129** |
| | | | | -10- -5s vs. 5s-10s | **0.0789** |
| **1U** | Repeated measures one-way ANOVA | n=4 mice, 2-4 trials per animal, total trials=12 | | Dunnet's multiple comparison test | |
| | F (1.470, 16.17) = 0.1879 | | **0.7635** | No significant differences | |
| **1V** | Repeated measures one-way ANOVA | n=4 mice, 5-6 trials per animal, total trials=21 | | Dunnet's multiple comparison test | |
| | F (2.100, 44.10) = 3.070 | | **0.0541** | -10- -5s vs. -5-0s | **0.0171** |
| | | | | -10- -5s vs. 0-5s | **0.7767** |
| | | | | -10- -5s vs. 5-10s | **0.9658** |
| **2C** | One-way ANOVA | n=3 mice, 9-23 trials each food, total trials=54 | **<0.0001** | Sidak's multiple comparison test | |
| | F (7, 100) = 15.01 | | | wood pre vs. wood post | **0.4519** |
| | | | | chow pre vs. chow post | **0.0007** |
| | | | | fam pre vs. fam post | **<0.0001** |
| | | | | nov pre vs. nov post | **<0.0001** |
| | | | | wood post vs. chow post | **0.5067** |
| | | | | fam post vs. wood post | **0.198** |
| | | | | nov post vs. wood post | **0.003** |
| | | | | fam post vs. chow post | **>0.9999** |
| | | | | nov post vs. chow post | **0.2472** |
| | | | | fam post vs. nov post | **0.2847** |
| **2D** | Repeated measures one-way ANOVA | n=3 mice | **0.0412** | Tukey's multiple comparison test | |
| | F (1.011, 2.022) = 22.29 | | | Chow vs. Familiar palatable food | **0.0339** |
| | | | | Chow vs. Novel palatable food | **0.0423** |
| | | | | Familiar vs. Novel palatable food | **0.1631** |
| **2E** | Repeated measures one-way ANOVA | n=3 mice | **0.038** | Tukey's multiple comparison test | |
| | F (1.005, 2.009) = 24.58 | | | Chow vs. Familiar palatable food | **0.0244** |
| | | | | Chow vs. Novel palatable food | **0.0443** |
| | | | | Familiar vs. Novel palatable food | **0.1238** |

**Table S2. Summary of statistical tests; animal studies**

| Figure | Statisitical test | n | p-value | Post test | p-value |
|---|---|---|---|---|---|
| **3C** | Two-tailed unpaired t-test (t=3.513, df=11) | Control=6; mPFC-LH hM4Di=7 | **0.0049** | | |
| **3G** | Two-way ANOVA | Control=7; mPFC-LH hM4Di=5 | | Sidak's multiple comparison test | |
| | Main effect of group (F (1, 10) = 0.6443) | | **0.4408** | No significant differences | |
| | Main effect of time (F (2, 20) = 28.48) | | **<0.0001** | | |
| | Interaction group*time (F (2, 20) = 0.4994) | | **0.6142** | | |
| **3H** | Two-way ANOVA | Control=11; mPFC-LH hM4Di=7 | | Sidak's multiple comparison test | |
| | Main effect of group (F (1, 16) = 0.06710) | | **0.3862** | No significant differences | |
| | Main effect of time (F (2, 32) = 291.8) | | **0.0001** | | |
| | Interaction group*time (F (2, 32) = 0.9802) | | **0.7989** | | |
| **3I** | Two-way ANOVA | Control=7; mPFC-LH hM4Di=5 | | Sidak's multiple comparison test | |
| | Main effect of group (F (1, 10) = 0.3343) | | **0.576** | No significant differences | |
| | Main effect of time (F (2, 20) = 142.4) | | **<0.0001** | | |
| | Interaction group*time (F (2, 20) = 1.077) | | **0.3596** | | |
| **3K** | Two-way ANOVA | Control=6; mPFC-LH hM4Di=7 | | Sidak's multiple comparison test | |
| **Fed** | Main effect of group (F (1, 11) = 0.02220) | | **0.8843** | No significant differences | |
| | Main effect of time (F (299, 3289) = 26.34) | | **<0.0001** | | |
| | Interaction group*time (F (299, 3289) = 0.3286) | | **>0.9999** | | |
| **3K** | Two-way ANOVA | Control=9; mPFC-LH hM4Di=7 | | Sidak's multiple comparison test | |
| **Fasted** | Main effect of group (F(1, 14) = 0.01138) | | **0.9166** | No significant differences | |
| | Main effect of time (F (10, 110) = 43.89) | | **<0.0001** | | |
| | Interaction group*time (F (299, 4186) = 0.2140) | | **>0.9999** | | |
| **3M** | Two-way ANOVA | Control=7; mPFC-LH hM4Di=6 | | Sidak's multiple comparison test | |
| **Home cage** | Main effect of group (F (1, 11) = 0.09474) | | **0.764** | No significant differences | |
| | Main effect of time (F (10, 110) = 25.63) | | **<0.0001** | | |
| | Interaction group*time (F (10, 110) = 0.1353) | | **0.9992** | | |

**Table S2. Summary of statistical tests; animal studies**

| Figure | Statisitical test | n | p-value | Post test | p-value |
|---|---|---|---|---|---|
| **3M** | Two-way ANOVA | Control=7; mPFC-LH hM4Di=6 | | Sidak's multiple comparison test | |
| **Novel cage** | Main effect of group (F (1, 11) = 2.693) | | **0.129** | 0 minutes | **>0.9999** |
| | Main effect of time (F (10, 110) = 14.52) | | **<0.0001** | 0.5 minutes | **>0.9999** |
| | Interaction group*time (F (10, 110) = 2.223) | | **0.0213** | 1 minute | **0.9925** |
| | | | | 1.5 minutes | **0.9874** |
| | | | | 2 minutes | **0.9998** |
| | | | | 2.5 minutes | **0.9693** |
| | | | | 3 minutes | **0.9693** |
| | | | | 3.5 minutes | **0.9693** |
| | | | | 4 minutes | **0.1829** |
| | | | | 4.5 minutes | **0.0132** |
| | | | | 5 minutes | **0.2765** |
| **3O** | Two-tailed unparied t-test (t=2.166, df=16) | Control=10; mPFC-LH hM4Di=8 | **0.0457** | | |
| **3P** | Two-tailed unparied t-test (t=2.188, df=16) | Control=10; mPFC-LH hM4Di=8 | **0.0439** | | |
| **3Q** | Two-tailed unparied t-test (t=1.434, df=16) | Control=10; mPFC-LH hM4Di=8 | **0.1709** | | |
| **3R % time eating** | Two-tailed unpaired t-test (t=2.263, df=16) | Control=10; mPFC-LH hM4Di=8 | **0.0379** | | |
| **% time sniffing food** | Two-tailed unpaired t-test (t=0.1187, df=16) | Control=10; mPFC-LH hM4Di=8 | **0.907** | | |
| **% time approaching food** | Two-tailed unpaired t-test (t=0.1187, df=16) | Control=10; mPFC-LH hM4Di=8 | **0.7842** | | |
| **% time away from food** | Two-tailed unpaired t-test (t=1.923, df=16) | Control=10; mPFC-LH hM4Di=8 | **0.0724** | | |
| **3T** | Two-tailed unpaired t-test (t=2.145, df=16) | Control=10; mPFC-LH hM4Di=8 | **0.0477** | | |
| **3U** | Two-tailed unpaired t-test (t=1.015, df=16) | Control=10; mPFC-LH hM4Di=8 | **0.3254** | | |
| **3V** | Two-tailed unpaired t-test (t=2.752, df=16) | Control=10; mPFC-LH hM4Di=8 | **0.0142** | | |
| **3W % time eating** | Two-tailed unpaired t-test (t=2.590, df=16) | Control=10; mPFC-LH hM4Di=8 | **0.0197** | | |
| **% time sniffing food** | Two-tailed unpaired t-test (t=0.5013, df=16) | Control=10; mPFC-LH hM4Di=8 | **0.623** | | |
| **% time approaching food** | Two-tailed unpaired t-test (t=0.7870, df=16) | Control=10; mPFC-LH hM4Di=8 | **0.4428** | | |
| **% time away from food** | Two-tailed unpaired t-test (t=2.656, df=16) | Control=10; mPFC-LH hM4Di=8 | **0.0173** | | |
| **4C** | Two-tailed unpaired t-test (t=3.541, df=14) | Control=9; mPFC-LH hM3Dq=7 | **0.0033** | | |
| **4G** | Two-way ANOVA | Control=11; mPFC-LH hM3Dq=9 | | Sidak's multiple comparison test | |
| | Main effect of group (F (1, 18)=5.202) | | **0.035** | 0 hours Control vs mPFC-LH hM3Dq | **>0.9999** |
| | Main effect of time (F (2, 36)=14.19) | | **<0.0001** | 2 hours Control vs mPFC-LH hM3Dq | **0.2718** |
| | Interaction group*time (F (2, 36)=5.510) | | **0.0082** | 4 hours Control vs mPFC-LH hM3Dq | **0.0019** |

**Table S2. Summary of statistical tests; animal studies**

| Figure | Statisitical test | n | p-value | Post test | p-value |
|---|---|---|---|---|---|
| **4H** | Two-way ANOVA | Control=11; mPFC-LH hM3Dq=9 | | Sidak's multiple comparison test | |
| | Main effect of group (F (1, 18)=2.775) | | **0.113** | 0 hours Control vs mPFC-LH hM3Dq | **>0.9999** |
| | Main effect of time (F (2, 36)=38.51) | | **<0.0001** | 2 hours Control vs mPFC-LH hM3Dq | **0.501** |
| | Interaction group*time (F (2, 36)=2.418) | | **0.1034** | 4 hours Control vs mPFC-LH hM3Dq | **0.0461** |
| **4I** | Two-way ANOVA | Control=9; mPFC-LH hM3Dq=7 | | Sidak's multiple comparison test | |
| | Main effect of group (F (1, 14)=3.987) | | **0.0657** | 0 hours Control vs mPFC-LH hM3Dq | **>0.9999** |
| | Main effect of time (F (2, 28)=139.5) | | **<0.0001** | 2 hours Control vs mPFC-LH hM3Dq | **0.3059** |
| | Interaction group*time (F (2, 28)=2.586) | | **0.0932** | 4 hours Control vs mPFC-LH hM3Dq | **0.0287** |
| **4L** | Two-way ANOVA | Control=8; mPFC-LH hM3Dq=7 | | Sidak's multiple comparison test | |
| | Main effect of group (F (1, 13)=3.556) | | **0.0819** | Fed Control vs mPFC-LH hM3Dq | **0.4697** |
| | Main effect of time (F (1, 13)=8.055) | | **0.014** | Fasted Control vs mPFC-LH hM3Dq | **0.0004** |
| | Interaction group*time (F (1, 13)=28.75) | | **0.0001** | | |
| **4M** | Two-way ANOVA | Control=8; mPFC-LH hM3Dq=7 | | Sidak's multiple comparison test | |
| | Main effect of group (F (1, 13)=0.5063) | | **0.4893** | No significant differences | |
| | Main effect of time (F (299, 3887)=41.38) | | **<0.0001** | | |
| | Interaction group*time (F(299, 3887)=3.361) | | **<0.0001** | | |
| **4N** | Two-way ANOVA | Control=8; mPFC-LH hM3Dq=7 | | Sidak's multiple comparison test | |
| | Main effect of group (F (1, 13)=5.628) | | **0.0338** | 4 hours 1 minute | **not significant** |
| | Main effect of time (F (299, 3887)=38.15) | | **<0.0001** | 4 hours 2 minutes | **0.0208** |
| | Interaction group*time (F (299, 3887)=10.54) | | **<0.0001** | 4 hours 3 minutes | **0.0208** |
| | | | | 4 hours 4 minutes | **0.0138** |
| | | | | 4 hours 5 minutes to 8 minutes | **0.0064** |
| | | | | 4 hours 9 minutes | **0.0024** |
| | | | | 4 hours 10 minutes to 24 minutes | **0.0021** |
| | | | | 4 hours 25 minutes to 26 minutes | **0.0017** |
| | | | | 4 hours 27 minutes | **0.0009** |
| | | | | 4 hours 28 minutes to 38 minutes | **0.0003** |
| | | | | 4 hours 39 minutes to 5 hours | **<0.0001** |
| **5E** | Two-tailed paired t-test (t=4.462, df=17) | n=4 mice, n=3-5 trials per animal, no. of pairs=18 | **0.0003** | | |

**Table S2. Summary of statistical tests; animal studies**

| Figure | Statisitical test | n | p-value | Post test | p-value |
|---|---|---|---|---|---|
| **5H** | Two-tailed paired t-test (t=5.274, df=14) | n=4 mice, n=2-4 trials per animal, no. of pairs=15 | **0.0001** | | |
| **5K** | One-way ANOVA | n=4 mice, 8-17 trials each food, total trials=47 | <0.0001 | Sidak's multiple comparison test | |
| | F (7, 86) = 6.382 | | | wood pre vs. wood post | **0.1316** |
| | | | | chow pre vs. chow post | **0.0337** |
| | | | | fam pre vs. fam post | **0.0011** |
| | | | | nov pre vs. nov post | **0.0037** |
| | | | | wood post vs. chow post | **>0.9999** |
| | | | | fam post vs. wood post | **0.9913** |
| | | | | nov post vs. wood post | **>0.9999** |
| | | | | fam post vs. chow post | **0.9968** |
| | | | | nov post vs. chow post | **>0.9999** |
| | | | | fam post vs. nov post | **0.9994** |

**Table S2. Summary of statistical tests; animal studies**

| Figure | Statisitical test | n | p-value | Post test | p-value |
| --- | --- | --- | --- | --- | --- |
| **6C** | Two-way ANOVA | WT=7; Vglut1-cre=15 | | Sidak's multiple comparison test | |
| | Main effect of group (F (1, 20) = 5.560) | | **0.0287** | 0 hours WT vs Vglut1-cre | **>0.9999** |
| | Main effect of time (F (2, 40) = 229.0) | | **<0.0001** | 1 hours WT vs Vglut1-cre | **0.0224** |
| | Interaction group *time (F (2, 40) = 4.968) | | **0.0118** | 2 hours WT vs Vglut1-cre | **0.0157** |
| **6E** | Two-tailed unpaired t-test (t=2.899, df=15) | WT=5; Vglut1-cre=12 | **0.011** | | |
| **6F** | Two-way ANOVA | | | Sidak's multiple comparison test | |
| | Main effect of group (F (1, 17) = 1.970) | | **0.1785** | WT No stim vs Stim | **0.6599** |
| | Main effect of stimulation (F (1, 17) = 1.615) | | **0.2209** | Vglut1-cre No stim vs Stim | **0.0056** |
| | Interaction group*stimuation (F(1, 17 =7.011)) | | **0.0169** | | |
| **S2G** | Two-tailed unpaired t-test (t=3.155, df=5) | n=6 mice, 1 2nd entry, 1 final entry per animal | **0.0252** | | |
| **S2I** | Two-tailed unpaired t-test (t=2.457, df=18) | n=10 mice | **0.0244** | | |
| **S2J** | Two-tailed unpaired t-test (t=2.457, df=18) | n=10 mice | **0.0153** | | |
| **S2K** | Two-tailed unpaired t-test (t=3.731, df=16) | n=9 mice | **0.0018** | | |
| **S3C** | Repeated measures one-way ANOVA | n=3 mice | | Tukey's multiple comparison test | |
| | F(1.179, 2.358) = 27.44 | | **0.0239** | Chow vs. Familiar palatable food | **0.1213** |
| | | | | Chow vs. Novel palatable food | **0.0447** |
| | | | | Familiar vs. Novel palatable food | **0.0854** |
| **S3D** | Two-tailed unpaired t-test (t=0.4601, df=17) | n=9 trials fed; n=10 trials fasted, 3 mice | **0.6513** | | |
| **S3E** | Two-tailed unpaired t-test (t=0.9339, df=20) | n=13 trials fed; n=9 trials fasted, 3 mice | **0.3615** | | |
| **S3F** | Two-tailed unpaired t-test (t=0.4620, df=31) | n=23 trials fed; n=10 trials fasted, 3 mice | **0.6473** | | |
| **S3G** | Two-tailed unpaired t-test (t=1.212, df=16) | n=9 trials fed; n=9 trials fasted, 3 mice | **0.243** | | |
| **S4B** | Two-way ANOVA | Control=5; mPFC-LH hM4Di=6 | | Sidak's multiple comparison test | |
| | Main effect of group (F(1, 9)=2.409)) | | **0.155** | No significant differences | |
| | Main effect of time (F(3.595, 32.36)=3.387)) | | **0.0234** | | |
| | Interaction group*time (F(5, 45)=1.485) | | **0.2135** | | |
| **S4C** | Two-tailed unpaired t-test (t=1.552, df=9) | Control=5; mPFC-LH hM4Di=6 | **0.155** | | |
| **S4D** | Two-way ANOVA | Control=5; mPFC-LH hM4Di=6 | | Sidak's multiple comparison test | |
| | Main effect of group (F(1, 9)=0.4027)) | | **0.5415** | No significant differences | |
| | Main effect of time (F(3.265, 29.38)=1.195)) | | **0.3306** | | |
| | Interaction group*time (F(5, 45)=1.998)) | | **0.0972** | | |
| **S4E** | Two-tailed unpaired t-test (t=0.6346, df=9) | Control=5; mPFC-LH hM4Di=6 | **0.5415** | | |

**Table S2. Summary of statistical tests; animal studies**

| Figure | Statisitical test | n | p-value | Post test | p-value |
|---|---|---|---|---|---|
| **S4F** | Two-way ANOVA | Control=7; mPFC-LH hM4Di=5 | | Sidak's multiple comparison test | |
| | Main effect of group (F(1, 10)=0.1348) | | **0.7211** | No significant differences | |
| | Main effect of time (F(3.669, 36.69)=2.493) | | **0.0644** | | |
| | Interaction group*time (F(5, 50)=1.806) | | **0.1287** | | |
| **S4G** | Two-tailed unpaired t-test (t=0.4360, df=10) | Control=7; mPFC-LH hM4Di=5 | **0.6721** | | |
| **S4H** | Two-way ANOVA | Control=7; mPFC-LH hM4Di=5 | | Sidak's multiple comparison test | |
| | Main effect of group (F(1,10)=0.2969) | | **0.5978** | No significant differences | |
| | Main effect of time (F(2.860, 28.60)=0.7017) | | **0.5524** | | |
| | Interaction group*time (F(5, 50)=1.718) | | **0.1476** | | |
| **S4I** | Two-tailed unpaired t-test (t=0.3904, df=10) | Control=7; mPFC-LH hM4Di=5 | **0.7044** | | |
| **S4J** | Two-tailed unpaired t-test (t=1.440, df=12) | Control=7; mPFC-LH hM4Di=7 | **0.1754** | | |
| **S4K** | Two-tailed unpaired t-test (t=1.581, df=12) | Control=7; mPFC-LH hM4Di=7 | **0.1398** | | |
| **S4L** | Two-tailed unpaired t-test (t=1.581, df=12) | Control=7; mPFC-LH hM4Di=7 | **0.1398** | | |
| **S4M % time grooming** | Two-tailed unpaired t-test (t=0.8254, df=12) | Control=7; mPFC-LH hM4Di=7 | **0.4252** | | |
| **% time rearing** | Two-tailed unpaired t-test (t=5.276, df=12) | Control=7; mPFC-LH hM4Di=7 | **0.0002** | | |
| **% time walking** | Two-tailed unpaired t-test (t=0.4186, df=12) | Control=7; mPFC-LH hM4Di=7 | **0.6829** | | |
| **% time stationary** | Two-tailed unpaired t-test (t=0.9389, df=12) | Control=7; mPFC-LH hM4Di=7 | **0.3663** | | |
| **S5B** | Two-way ANOVA | Control=12; mPFC-LH hM3Dq=9 | | Sidak's multiple comparison test | |
| | Main effect of group (F(1, 19) 0.05376) | | **0.8191** | No significant differences | |
| | Main effect of time (F(3.678, 69.89)=0.8761) | | **0.4756** | | |
| | Interaction group*time (F(5, 95)=2.206) | | **0.06** | | |
| **S5C** | Two-tailed unpaired t-test (t=0.1763, df=19) | Control=12; mPFC-LH hM3Dq=9 | **0.8619** | | |
| **S5D** | Two-way ANOVA | Control=12; mPFC-LH hM3Dq=9 | | Sidak's multiple comparison test | |
| | Main effect of group (F(1, 19)=0.6096) | | **0.4446** | No significant differences | |
| | Main effect of time (F(3.976, 75.54)=1.304) | | **0.2761** | | |
| | Interaction group*time (F(5, 95)=1.343) | | **0.2532** | | |
| **S5E** | Two-tailed unpaired t-test (t=0.8698, df=19) | Control=12; mPFC-LH hM3Dq=9 | **0.3953** | | |
| **S5F** | Two-way ANOVA | Control=11; mPFC-LH hM3Dq=8 | | Sidak's multiple comparison test | |
| | Main effect of group (F(1, 17)=0.4503) | | **0.5112** | No significant differences | |
| | Main effect of time (F(3.719, 63.23)=3.641) | | **0.0115** | | |
| | Interaction group*time (F(5, 85)=0.8161) | | **0.5415** | | |

**Table S2. Summary of statistical tests; animal studies**

| Figure | Statisitical test | n | p-value | Post test | p-value |
|---|---|---|---|---|---|
| **S5G** | Two-tailed unpaired t-test (t=0.5888, df=17) | Control=11; mPFC-LH hM3Dq=8 | **0.5637** | | |
| **S5H** | Two-tailed unpaired t-test (t=1.183, df=18) | Control=11; mPFC-LH hM3Dq=8 | | | |
| | Main effect of group (F(17, 85)=1.840) | | **0.3647** | | |
| | Main effect of time (F(3.576, 60.79)=1.574) | | **0.1978** | | |
| | Interaction group*time (F(5, 85)=0.5655) | | **0.7262** | | |
| **S5I** | Two-tailed unpaired t-test (t=0.9313, df=17) | Control=11; mPFC-LH hM3Dq=8 | **0.3647** | | |
| **S5J** | Two-tailed unpaired t-test (t=0.3745, df=18) | Control=11; mPFC-LH hM3Dq=9 | **0.7124** | | |
| **S5K** | Two-tailed unpaired t-test (t=1.183, df=18) | Control=11; mPFC-LH hM3Dq=9 | **0.252** | | |
| **S5L** | Two-tailed unpaired t-test (t=1.037, df=18) | Control=11; mPFC-LH hM3Dq=9 | **0.3133** | | |
| **S5M % time rearing** | Two-tailed unpaired t-test (t=0.03213, df=18) | Control=11; mPFC-LH hM3Dq=9 | **0.9747** | | |
| **% time walking** | Two-tailed unpaired t-test (t=1.140, df=18) | Control=11; mPFC-LH hM3Dq=9 | **0.2691** | | |
| **% time stationary** | Two-tailed unpaired t-test (t=1.003, df=18) | Control=11; mPFC-LH hM3Dq=9 | **0.3292** | | |
| **S6E** | Two-way ANOVA | WT=5; Vglut1-cre=12 | | Sidak's multiple comparison test | |
| | Main effect of group (F (1, 15) = 1.678) | | **0.2148** | No significant differences | |
| | Main effect of time (F (1, 15) = 4.787) | | **0.0449** | | |
| | Interaction group*time (F (1, 15) = 0.1804) | | **0.6771** | | |
